# Changes in the distribution of fitness effects and adaptive mutational spectra following a single first step towards adaptation

**DOI:** 10.1101/2020.06.12.148833

**Authors:** Dimitra Aggeli, Yuping Li, Gavin Sherlock

## Abstract

The fitness effects of random mutations are contingent upon the genetic and environmental contexts in which they occur, and this contributes to the unpredictability of evolutionary outcomes at the molecular level. Despite this unpredictability, the rate of adaptation in homogeneous environments tends to decrease over evolutionary time, due to diminishing returns epistasis, causing relative fitness gains to be predictable over the long term. Here, we studied the extent of diminishing returns epistasis and the changes in the adaptive mutational spectra after yeast populations have already taken their first adaptive mutational step. We used three distinct adaptive clones that arose under identical conditions from a common ancestor, from which they diverge by a single point mutation, to found populations that we further evolved. We followed the evolutionary dynamics of these populations by lineage tracking and determined adaptive outcomes using fitness assays and whole genome sequencing. We found compelling evidence for diminishing returns: fitness gains during the 2^nd^ step of adaptation are smaller than those of the 1^st^ step, due to a compressed distribution of fitness effects in the 2^nd^ step. We also found strong evidence for historical contingency at the genic level: the beneficial mutational spectra of the 2^nd^-step adapted genotypes differ with respect to their ancestor and to each other, despite the fact that the three founders’ 1^st^-step mutations provided their fitness gains due to similar phenotypic improvements. While some targets of selection in the second step are shared with those seen in the common ancestor, other targets appear to be contingent on the specific first step mutation, with more phenotypically similar founding clones having more similar adaptive mutational spectra. Finally, we found that disruptive mutations, such as nonsense and frameshift, were much more common in the first step of adaptation, contributing an additional way that both diminishing returns and historical contingency are evident during 2^nd^ step adaptation.

## Introduction

Stephen Jay Gould argued that historical contingency makes evolutionary outcomes largely unpredictable, and that were we to replay the “tape of life”, we would likely end up with a different world each time^1^. However, frequently observed instances of both parallel^2–4^ and convergent^5–7^ evolution suggest that, at least under some circumstances, adapting populations may simply take different paths to the same peak on a fitness landscape. Environmental similarities, genotypic relatedness and proximity to an optimum in the fitness landscape constitute some of the constraints contributing to convergent or parallel adaptive responses^4,8–21^.

Closely related genotypes are often employed to study the effects of evolutionary history on adaptation in various experimental systems^22–30^. A frequent observation is that fitness gains decrease over time during adaptive evolution—termed diminishing returns—most convincingly demonstrated in cases where founders with differing initial fitness are used^24,25,27,28,30^. However, support for the role of historical contingency during adaptation is not uniformly consistent. For example, evolutionary history has been shown to both contribute^23^ and not contribute^26^ to defining subsequent adaptive mutational spectra in closely related *Pseudomonas aeruginosa* lineages, while historical contingency in related evolving *Escherichia coli* populations manifested at a phenotypic but not at a molecular level^29^. By contrast, evidence of first-step adaptive mutations in *Saccharomyces cerevisiae* being mutually exclusive due to reciprocal sign epistasis^22,31,32^ is clearly supportive of historical contingency. Nevertheless, experiments founded with related *S. cerevisiae* clones spanning a range of fitness effects, suggest that convergence at a molecular level can and does occur^30^. Such apparently contradictory results may stem from differences in evolutionary timescales, population sizes and culture conditions (serial transfer vs. continuous culture in chemostats). For example, longer timescales (up to several hundreds of generation) that allow for the rise of lineages to frequencies sufficient for easy detection via sequencing, also result in clonal interference, a consequence of clonal propagation in well-mixed environments^24,33–37^. Given enough time, clonal interference will result in a somewhat predictable outcome because of competition among adaptive lineages that will reproducibly lead to fixation of those with the highest fitness^30,34,38–40^.

The application of molecular barcoding to experimental microbial evolution (EME), for the purpose of tracking lineages, has enabled high-resolution characterization of evolutionary processes^4,20,21,39–41^, importantly, on shorter timescales (less than a few hundred generations). Such studies have revealed a plethora of available adaptive mutations that increase in frequency early in the evolution, but most of which will eventually go extinct due to being outcompeted by high fitness lineages later in the evolution^4,39,41^. The increase in our detection limit permitted by lineage tracking has allowed high resolution characterization of these adaptive events that typically go extinct in experimental evolutions. This has provided us the opportunity to re-examine the prevalence of diminishing returns and historical contingency, while taking into consideration both evolutionary timescales and fitness effects.

Here we used DNA barcoding to investigate how closely-related adapted genotypes of *S. cerevisiae*, each with a single mutation relative to their common ancestor, evolve in the environment under which they were originally selected^4,41^. The evolutionary environment is serial-transfer under glucose-limitation, where cells undergo lag, fermentation and respiration growth phases within each growth cycle. Common adaptive strategies of the 1^st^ step adaptation in this environment included upregulation of the RAS/PKA and TOR/Sch9 pathways^4^; our founders for the 2^nd^-step evolutions carry either a *cyr1*, a *gpb2* (both of which upregulate the Ras/PKA pathway), or a *tor1* (which upregulates the TOR/Sch9 pathway) mutation. All three of these mutants have increased cell size and higher fitness relative to their ancestor^4,20^, though their fitness advantages manifest differently within lag, fermentation and respiration growth phases^20^. We evolved barcoded populations of each of these three mutants and characterized rates of adaptation, and the distributions of fitness effects (DFE) of second step mutations. We then isolated hundreds of independent adaptive lineages and performed whole genome sequencing and fitness remeasurements. We found that 2^nd^-step mutations confer a smaller fitness advantage than the 1^st^-step mutations in the respective backgrounds where they arise, consistent with diminishing returns epistasis. We also found that there is a partial overlap in the molecular basis of the 2^nd^-step of adaptation between genetic backgrounds: the TOR/Sch9 pathway mutant frequently adapted via mutations in the RAS/PKA pathway, while the RAS/PKA pathway mutants, *cyr1* and *gpb2*, sometimes acquired mutations in the TOR/Sch9 pathway. On the other hand, we rarely identify second-step mutations that further modify the same pathway. We also found that the spectrum of adaptive mutations shifted from affecting pathways that regulate the cell cycle and nutrient signaling to pathways that affect stress responses. Targets of selection include genes in the HOG, retrograde flow (RTG) and glutathione biosynthesis pathways. Whereas *GSH1*, which functions in the glutathione biosynthetic pathway, was a target of selection in all backgrounds, the HOG pathway was targeted only in the TOR/Sch9 pathway mutant and the RTG pathway was mutated in the RAS/PKA mutants. The RAS/PKA pathway mutants had similar relative changes in fitness and adaptive mutational spectra to one another, that differed from those of the TOR/Sch9 pathway mutant. Finally, we found that the second step mutations were less likely to be disruptive (nonsense and frameshift mutations) compared to first step mutations. Altogether, our data show that a single adaptive change is sufficient to cause further genetic divergence during adaptation, demonstrating that historical contingency does influence the outcome evolutionary outcomes, and that the DFE between first and second step mutations differs, consistent with diminishing returns epistasis.

## Materials and Methods

### Strains and strain handling

All strains used are S288C derivatives, which were evolved and characterized previously^4,20,41^ (Table 1). Yeast strains and pools were saved as glycerol stocks at −80°C. Yeast transformations were performed by a lithium acetate/PEG method^42^.

**Table 1.** Strains used in this study

| Strain | Genotype | Description | Reference |
| --- | --- | --- | --- |
| GSY5096/<br>SHA118 | *MATalpha, ura3Δ, ybr209w::GalCre-natMX* | wild-type universal ancestor | [35] |
| GSY5375 | *MATa, ura3Δ, ybr209w::GalCre-natMX* | MATa version of the ancestor, used for back-crossing | this study |
| GSY6701 | *MATalpha, ura3Δ, ybr209w::GalCre-natMX, cyr1<sup>S917Y</sup>* | evolution founder | this study |
| GSY6702 | *MATalpha, ura3Δ, ybr209w::GalCre-natMX, gpb2<sup>Y282\*</sup>* | evolution founder | this study |
| GSY6703 | *MATalpha, ura3Δ, ybr209w::GalCre-natMX, tor1<sup>F1712L</sup>* | evolution founder | this study |
| GSY5481 | *MATalpha, ura3Δ, ybr209w::full barcode, cyr1<sup>S917Y</sup>* | evolved clone | [4, 35] |
| GSY5128 | *MATalpha, ura3Δ, ybr209w::full barcode, gpb2<sup>Y282\*</sup>* | evolved clone | [4, 35] |
| GSY5153 | *MATalpha, ura3Δ, ybr209w::full barcode, tor1<sup>F1712L</sup>* | evolved clone | [4, 35] |

### Yeast Growth Media and Growth Cycle

Evolutionary and fitness remeasurement conditions matched those used previously^4,41^. Briefly, M3 medium, consisting of 5x Delft medium^43^ with 4% ammonium sulfate and 1.5% dextrose, was used. Serial batch cultures were conducted by growing cells in 100 mL M3 medium in 500 mL Delong flasks (Bellco) at 30°C and 223 RPM. Yeast was grown for 48 hours between transfers and for each new cycle 400 μL of the grown culture (~8 × 10^7^ cells) were used as inoculum for the new culture, resulting in a 1:250 bottleneck.

### Construction and characterization of the founder strains of the evolutions

The prelanding pad strain (SHA118, Table 1) that is receptive to barcoding^41^ was transformed with a galactose-inducible HO-containing plasmid. The strain diploidized upon exposure to galactose and a diploid clone was sporulated and dissected; a *MATa* derivative was isolated (GSY5375, Table 1) and saved for subsequent crosses with the evolved clones. Loss of the HO-containing plasmid was verified by absence of growth on appropriate selective medium. Strains GSY5481, GSY5128 and GSY5153, derived from evolution under glucose limitation and previously characterized^4,20,41^ (Table 1), were backcrossed twice to GSY5375. Competitive fitness of segregants and evolved parents was assayed in triplicate compared to a fluorescent derivative of the ancestor, as in ^4^, with the following modifications: to increase throughput the assays were performed in 5 mL cultures in tubes incubated in a roller drum, instead of 100 mL cultures in flasks. As a result, the fitness estimates deviate from those previously reported^4^ (Fig. S1, Table S1). Additionally, since the derived strains do not contain a barcode (which reconstitutes a *URA3* gene), they require uracil, so the fitness assays were performed in M3 medium supplemented with uracil. All segregants were genotyped for the variant of interest by amplification of the respective locus and Sanger sequencing. The oligos used for genotyping are shown in table S2.

### Barcoding

Strains GSY6701, GSY6702 and GSY6703 (Table 1) were transformed with a low and high complexity barcode, consecutively. These strains have the *YBR209w* locus replaced with the prelanding pad (corresponding to strain SHA118^41^). The low complexity barcode was derived by PCR amplification of part of the L001 plasmid library, containing the lox66 site, the DNA barcode, the artificial intron, the 3’ half of *URA3*, and *HygMX*^44^. The fragment was amplified with primers BC_F-DY and BC_R1-DY (Table S2), from 12 ng of L001 library in a 50 μL reaction with PrimeSTAR (TAKARA, Mountain View, CA) using the following conditions: hot start, initial denaturation at 98°C for 2’, 30 cycles of 98°C for 30”, 55°C for 15” and 72°C for 3’, and final extension at 72°C for 10’. The product was purified with the QIAquick PCR purification kit (QIAGEN, Germantown MD), transformed into each of GSY6701, GSY6702 and GSY6703 and successful transformants were selected on YPD + Hygromycin. Single transformants were further transformed with the high complexity library (pBAR3)^41^ with the following modification: after transformation the cells were grown on liquid YP + 2% galactose for ~16 hours for Cre recombinase induction prior to selection on SC-ura plates with 2% glucose. Cell growth was estimated by cell counting immediately after transformation and before plating. The number of unique transformants was estimated by plating a dilution on selective medium and correcting for growth. After one day of growth on selective medium the transformants were pooled and saved as glycerol stocks at −80°C (high complexity subpools with a common low complexity barcode). The final founding pools for the evolutions were constructed by pooling high complexity subpools to an estimated total of ~700,000 unique transformants per initial clone.

### Evolution experiments

Evolution experiments were conducted as described previously^41^. Briefly, 10^8^ cells of each of the founding populations were used to inoculate 100 mL of SC-ura, 2% dextrose, supplemented with hygromycin in 500 mL Delong flasks (Bellco). The cells were grown for 24 hours at 30°C and 223 RPM, the end of which was considered generation 0 of the evolution experiment. 400 μL of the initial culture were used to inoculate M3 medium in duplicate, as described in the ‘Yeast Growth Media and Growth Cycle’ section. The evolution experiments were conducted for a total of 20 transfers, corresponding to approximately 160 generations. Prior to each transfer the medium was prewarmed at 30°C for 1 hour. For each timepoint, 2 × 1 mL aliquots were saved as glycerol stocks at −80°C and the rest of the culture was spun down, resuspended in 5 mL sorbitol buffer (0.9 M sorbitol, 100 mM Tris pH 7.5, 100 mM EDTA), aliquoted in Eppendorf tubes (~1 mL each), and saved at −20°C to be used for genomic DNA and barcode library preparations.

### Clone isolation

Individual clones were sorted at the Stanford Shared FACS facility either from all timepoints (one or two 96-well plates each) for ploidy determination, or from selected timepoints (10 plates each of the following timepoints: *cyr1* evolution, replicate 1, timepoint 20 (generation 160), *gpb2* evolution, replicates 1 and 2, timepoint 13 (generation 104) and *tor1* evolution, replicate 1, timepoint 12 (generation 96)) for fitness remeasurements, ploidy determination and whole genome sequencing.

### Ploidy assay

Ploidy was determined with a high-throughput benomyl-based assay as described^4^.

### Genomic DNA and library preparation for barcode lineage tracking

Genomic DNA was prepared as follows. An aliquot of cells stored at −20°C was thawed at room temperature. The cells were spun down, washed once in water, resuspended in 400 μL lysis buffer (0.9 M sorbitol, 50 mM Na phosphate pH 7.5, 240 μg/mL zymolase, 14 mM β-mercaptoethanol) and incubated at 37°C for 30 minutes. After the incubation, 40 μL 0.5 M EDTA, 40 μL 10% SDS and 56 μL 20 mg/mL proteinase K (Life Technologies 25530-015) were added consecutively, with brief vortexing after each addition, and the samples were incubated at 65°C for 30 minutes. Subsequently, the samples were cooled on ice for 5’, 200 μL of 5 M potassium acetate were added, and the samples were mixed by shaking and incubated on ice for an additional 30 minutes. Following incubation, the samples were spun down at full speed in a microcentrifuge for 10 minutes, and the supernatant was transferred to a new tube with 750 μL isopropanol and was let to rest on ice for 5 minutes. The precipitated nucleic acid was spun down full speed in a microcentrifuge for 10 minutes and washed twice with 70% ethanol. After the second wash the nucleic acid was let to dry completely and then it was resuspended in 50 μL 10 mM Tris pH 7.5. Overnight incubation at room temperature or short incubation at 65°C sometimes was necessary for complete resuspension. RNA was digested with the addition of 0.5 μL 20 mg/mL RNase A (Fisher Scientific, Waltham MA) and incubation at 65°C for 30 minutes.

A two-step PCR protocol was used to amplify the barcoded locus (see ^4^ for primer details). The first amplification was conducted using OneTaq 2X Master Mix (NEB, Ipswich MA), a total of 6 μg genomic DNA and a limited amount of primers in 6 × 50 μL reactions with the following composition: 1X OneTaq Mix, 50 nM each forward and reverse primer, 2 mM MgCl_2_, 20 ng/μL gDNA, in the following conditions: hot start, initial denaturation at 94°C for 10’, 3 cycles of 94°C for 3’, 55°C for 1’ and 68°C for 1’, and final extension at 68°C for 1’. The 6 reactions were combined, purified using the QIAquick PCR purification kit (QIAGEN, Germantown MD) and eluted into 30 μL EB buffer. All the eluate was used as template in a single 50 μL 2^nd^ reaction, with the following composition: 0.5 μL Herculase II fusion DNA polymerase (Agilent, Santa Clara CA) per 50 μL reaction, 1xHerculase buffer, 1 mM dNTPs and 500 nM each of PE1 and PE2^4^, and was amplified in the following conditions: hot start, initial denaturation at 98°C for 2’, 20 cycles of 98°C for 10”, 69°C for 20” and 72°C for 30”, and final extension at 72°C for 1’. Barcode libraries were pooled isostoichiometrically and sequenced on an Illumina NextSeq 550.

### DFE / Mutational Fitness Spectrum u(s) inference

Lineage tracking from barcode sequencing was reconstructed as described in ^41^ and using https://github.com/Sherlock-Lab/Barcode_seq/blob/master/bartender_BC1_BC2.py with some minor modifications. Briefly, after extraction of the UMI, and both low and high complexity barcodes from the sequencing read, low complexity barcodes were clustered against their expected sequences, whereas the high complexity barcodes were pooled across all libraries and clustered with bartender (v1.1)^45^. The updated reads and the UMIs were used to derive raw barcode counts, which were assembled into the raw count lineage trajectories. Low coverage timepoints and barcodes that appeared in a single timepoint (considering replicate evolutions) or had no reads at timepoint 0 were excluded from subsequent analysis. The included timepoints and the number of reads and barcodes per timepoint are shown in table S3. Filtered raw count lineage trajectories are provided for each replicate evolution (Supplemental data: lineage trajectory data).

Using the lineage frequency changes over time, lineages’ fitness per generation (s) and establishment time (tau) were estimated using the same method as in ^41^. Lineages with reads between 20-30 at each timepoint were treated as neutral and were used to estimate population mean fitness. Lineage tracking data from generation 0 to generation 136 were used for fitness inference in all evolutions, except for *gpb2* evolution replicate 1, for which we only had adequate data up to generation 120. Lineage tracking data for the ancestor up to generation 112 and 96, for replicates 1 and 2, respectively, were used for fitness inference as in ^41^. The generations chosen are the times at which adapted lineages have reached a sufficient frequency in the population, while the majority of such lineages theoretically carry a single beneficial mutation.

Mutations can occur during the barcoding process and prior to the onset of the experiment, some of which can be beneficial in the evolutionary condition^41^. To characterize the mutational rate during the evolution, lineages with such pre-existing mutations were removed from fitness inference. The following two criteria were used to define lineages with pre-existing mutations: 1) being adaptive in both evolutionary replicates and 2) having an establishment time < −2/s in at least one replicate.

Mutation rates in different fitness intervals were calculated using equation 101 in ^41^:

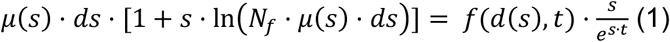

where *ds* = 0.002 is the fitness interval considered, *μ*(*s*) the mutation rate within a specific fitness interval [*s*, *s* + *ds*], *N_f_* = 10^12^ the approximated largest size the population has reached during the barcoding process, and *f*(*d*(*s*, *t*) the summed frequency of lineages whose fitness are within the interval [*s*, *s* + *ds*] at generation *t*. The error of the estimated *μ*(*s*) is 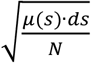, where *N* = 6 ∙ 10^8^, is the approximated effective population size during evolution.

Note that the barcode sequencing coverage of the ancestor evolutions was ~10-20X higher than those of *cyr1*, *gpb2* and *tor1* evolutions (Table S3, compare to Table 2 in Supplementary information in ^41^). To avoid biases introduced by sequencing coverage differences, we down-sampled the ancestor sequencing data to a depth comparable to those of the *cyr1*, *gpb2* and *tor1* evolutions: 2 × 10^7^ at time 0 and 3 × 10^6^ at the rest of the timepoints. Fitness was inferred before and after down-sampling. Lineages with 5-10 reads were treated as neutrals to infer the population mean fitness (vs 20-30 used in the full datasets).

**Table 2.** Mutation rates per genotype from mutational fitness spectra

| genotype | evolution | diploidization<br>fitness | density<br>(diploidization<br>rate) | diploidization<br>rate | average<br>diploidization<br>rate | high fitness<br>(>0.05)<br>mutation<br>rate | average<br>high fitness<br>(>0.05)<br>mutation<br>rate | high fitness<br>(0.07-0.12)<br>mutation<br>rate | high fitness<br>(>0.12)<br>mutation<br>rate |
| --- | --- | --- | --- | --- | --- | --- | --- | --- | --- |
| WT | evo1 | 0.032 | 1.53E-02 | 3.05E-05 | 2.03E-05 | 1.42E-06 | 5.32E-06 | 5.18E-07 | 8.84E-10 |
|  | evo2 | 0.041 | 5.03E-03 | 1.01E-05 |  | 9.22E-06 |  | 7.57E-07 | 2.27E-08 |
| *cyr1* | evo1 | 0.036 | 1.27E-04 | 2.54E-07 | 3.29E-07 | 7.74E-08 | 8.49E-08 | 6.92E-09 | 0.00E+00 |
|  | evo2 | 0.029 | 2.02E-04 | 4.04E-07 |  | 9.25E-08 |  | 6.38E-09 | 8.59E-13 |
| *gpb2* | evo1 | 0.029 | 6.62E-04 | 1.32E-06 | 1.19E-06 | 1.27E-07 | 9.56E-08 | 1.14E-08 | 5.59E-12 |
|  | evo2 | 0.034 | 5.25E-04 | 1.05E-06 |  | 6.41E-08 |  | 9.48E-09 | 7.69E-14 |
| *tor1* | evo1 | 0.036 | 8.45E-04 | 1.69E-06 | 3.35E-06 | 3.39E-07 | 2.01E-07 | 8.69E-09 | 0.00E+00 |
|  | evo2 | 0.030 | 2.50E-03 | 5.01E-06 |  | 6.23E-08 |  | 8.56E-10 | 0.00E+00 |

### Barcode determination of isolated clones

To identify the barcodes of the isolated clones in 96-well plates, we employed a 2- (column, row) or 3- (column, row, plate) dimensional pooling strategy, inspired by ^46^. Briefly, we arranged 20 plates per batch into a 4 columns × 5 rows plate matrix and constructed 48 column pools from clones out of 40 wells each and 40 row pools from clones out of 48 wells each. For the second batch we included semi-redundant half-plate pools (40 pools from clones out of 48 wells each) to increase the successful barcode recovery rate. We pooled our samples after cell growth and prepared barcode libraries for Illumina sequencing. Barcodes were recovered for each well at a rate of ~90%, which was somewhat dependent on the barcode diversity of the sampled timepoint (identical barcodes in multiple wells makes it more challenging algorithmically to match barcodes to wells).

### High-throughput Fitness Measurements and Analysis

#### Pooling of clones

Clones isolated from the evolutions were pooled together for high-throughput fitness assays. We used a multi-pronged pinner to take clones from frozen stock and pin them into a set of 96 deep-well plates with 700 μL YPD medium in each well. Cells were grown at 30°C for 2 days to reach saturation without shaking. 500 μL of 50% glycerol were added into each well using a multichannel pipette. 1 mL of the mixture from each well was pooled, and the final pool was mixed and aliquoted into 2 mL Eppendorf tubes, which were stored at −80°C for future fitness measurements.

#### Preculture

Each replicate fitness experiment was initiated with a 1 mL frozen aliquot of the pooled cell culture, thawed at room temperature, and inoculated into 15 mL M3 in a 500 mL Delong flask. The culture was grown at 30°C and 223 RPM overnight for cell propagation. 400 μL of the overnight culture were inoculated into 100 mL of fresh M3 medium and precultured at the standard condition for 2 days.

A derivative of the ancestor carrying a restriction site in the barcode region was used to compete with the pool of evolved clones for fitness measurements^4^. The ancestor clone was resurrected from frozen stock onto M3 agar plates and grown for 2 days until colonies were visible. A single colony was inoculated into 3 mL of M3 medium and grown for 48 hours (30°C in a roller drum). 400 μL of that culture were used to inoculate precultures (100 mL M3 medium in 500 mL Delong flasks, 223 RPM 30°C).

#### Competition

Fitness assays were conducted by mixing the pooled preculture with the ancestor preculture in a 1:9 ratio (time 0) and growing the resulting population for four successive growth cycles (timepoints 1, 2, 3 and 4), under the evolutionary condition. At the end of each cycle, 400 μL cell culture were inoculated into 100 mL fresh media to start the next cycle. Cells were collected at time 0, and at the end of each of the four growth cycles. The cell pellet from each sample was resuspended in 5 mL sorbitol solution (0.9 M sorbitol, 0.1 M Tris-HCL pH 7.5, 0.1 M EDTA pH 8.0), aliquoted into 2 mL Eppendorf tubes and stored at −20°C. Three technical replicates were performed per fitness assay. Genome extraction, barcode amplification and Illumina sequencing were conducted for each sample (timepoint and replicate).

#### Genomic DNA Sample Preparation

Genomic DNA was isolated and treated as described in the ‘Genomic DNA and library preparation for barcode lineage tracking’ section.

#### Fitness Estimation

DNA barcodes were sequenced on an Illumina NextSeq 500/550 platform and their abundances were used to estimate lineages’ frequencies in the population, as previously described^4^. Fitness estimates were conducted for all clones against the neutrals from the wild type evolution and for clones derived from each ancestral genotype separately against the neutrals of the specific genotype. The source code for computing these fitness estimates can be found at https://github.com/barcoding-bfa/fitness-assay-python. We ran two iterations of the script. First, we used all barcode counts as input and recovered fitness estimates and barcodes that were likely to be neutral. Barcodes identified by the first iteration were associated with their physical position on the 96-well plates in frozen stock, and the ploidy of the clones they represent. For the second iteration, apart from the barcode counts, a list of specifically haploid neutral clones was also provided (this is an optional argument of the fitness estimation algorithm). Fitness estimates from the 2^nd^ run were used for further analysis. Final fitness estimates were calculated by inverse variance weighting of estimates from all three replicates.

### Genome-wide sequencing library preparation

Genomic DNA libraries were constructed as described previously^40^. Briefly, clones selected for sequencing were grown in 500 μL YPD in 96 deep-well plates for two days at 30°C without shaking. 400 μL of saturated cell culture were used for DNA extraction with the Invitrogen PureLink Pro 96 Genomic DNA Kit (Catalog no. K1821-04A) in a 96-well format. Libraries were prepared and multiplexed with Nextera technology, and a high throughput protocol^47^. Samples were sequenced on an Illumina HiSeq 4000 with 2×150 bp paired end reads.

### Variant calling

SNP, small indel and structural variants were called for 105 clones using Sentieon Genomic Tools Version 201711.02, as described previously^21^. Briefly, FASTQ files were trimmed using cutadapt version 1.16^48^ and trimmed reads were mapped to the *S. cerevisiae S288C* reference genome R64-1-1 (https://downloads.yeastgenome.org/sequence/S288C_reference/genome_releases/) using bwa^49^. Mapped and sorted reads were then used for the variant calling. Variants were further annotated using snpEff and SNPSift^50^. The source code for variant calling and annotation can be found at https://github.com/liyuping927/DNAscope-variants-calling.

### Variant filtering

To eliminate false positive variants, we applied the following filters. First, variants from lineages with an average genome-wide coverage < 10, and all mitochondrial variants were filtered out. Second, variants in *FLO1* and *FLO9* genes were filtered out due to poor alignment in both genomic regions. Third, variants present in more than five clones and at least two genetic backgrounds out of *CYR1*, *GPB2* and *TOR1* mutants, they were likely present in the common ancestor and were filtered out. Fourth, variants with a quality score < 150 and only occurring in one clone were filtered out. Locus alignment against the reference genome was visually inspected to assess variants present in more than one clone, but with a quality score < 150 in at least one of them. Provided that the implicated loci were well-covered and not in highly repetitive regions, the variants were considered *bona fide* regardless of their quality score. Otherwise, they were discarded in all clones where they occurred. Lastly, we manually verified variants that passed the above filtering by inspecting the corresponding loci alignments against the reference genome and further filtering out false positives, typically occurring in highly repetitive or poorly sequenced regions.

## Results

### Experimental design

Previously, we evolved a population of barcoded haploid yeast cells in a 2-day serial transfer condition under glucose limitation and isolated thousands of evolved clones from cycle 11 (after ~88 generations)^4,41^. Such a timescale was long enough for adaptive clones to rise to a sufficient frequency in the population, while short enough that the majority of adaptive clones carries only a single causative mutation. We then measured the fitness of thousands of isolated clones under the evolutionary condition and whole genome sequenced hundreds of adaptive clones to identify their causative mutations^4^. Two major adaptive strategies were observed: self-diploidization, and upregulation of nutrient-sensing pathways, including the RAS/PKA pathway and the TOR/Sch9 pathway^4^. We refer to this prior evolution experiment as the “1^st^-step evolution”.

Here, we chose three adapted clones from the 1^st^-step evolution. Compared to their common ancestor, each adapted clone carries one of the following mutations: a presumptive gain-of-function mutation in a positive regulator (*cyr1*) of the Ras-PKA pathway, a loss-of-function mutation in a negative regulator (*gpb2*) of the same pathway, and a presumptive gain-of-function mutation in a positive regulator (*tor1*) of the Tor pathway (Table 1). The founding populations were derived from adapted clones via backcrossing with a mating-type switched version of the unbarcoded wild-type ancestor (strain GSY5375, Table 1). Fitness advantages of the derived strains were validated and shown to be monogenic and segregate with the mutation (Fig. S1, Table S1). These derivatives were then re-barcoded and further evolved for 160 generations in the same environment. We refer to this further evolution as the “2^nd^-step evolution” (Fig. 1). We performed low coverage barcode sequencing (average of 27 reads per barcode for timepoint 0 and 12 reads per barcode for the rest; see Table S3) of the populations over the course of the evolutions (Fig. S2) and used these data to estimate the fraction of adapted individuals at each timepoint (Fig. S3). Based on these data, as well as ploidy assays (Fig. S4), we isolated thousands of clones from cycles 20, 13 and 12, corresponding to generations 160, 104 and 96 (cells roughly divide 8 times during each cycle), from the “2^nd^-step evolution” of *cyr1*, *gpb2* and *tor1*, respectively, where ~25% to 50% of the individuals in the population are estimated to be adaptive. Fitness remeasurements and genome-wide sequencing were conducted for these evolved clones isolated from the 2^nd^ step evolution. Fitness estimates are expressed per generation (assuming 8 generations per growth cycle) for consistency with the bulk of the literature, although we are aware that fitness advantage is not equally distributed within the growth cycle^20^. In this study, we refer to the original ancestor used in the 1^st^-step evolution as the “wild-type” ancestor and we refer to the founders of the 2^nd^-step evolutions, *tor1*, *gpb2* and *cyr1* mutants, as “adapted” ancestors.

**Figure 1.**
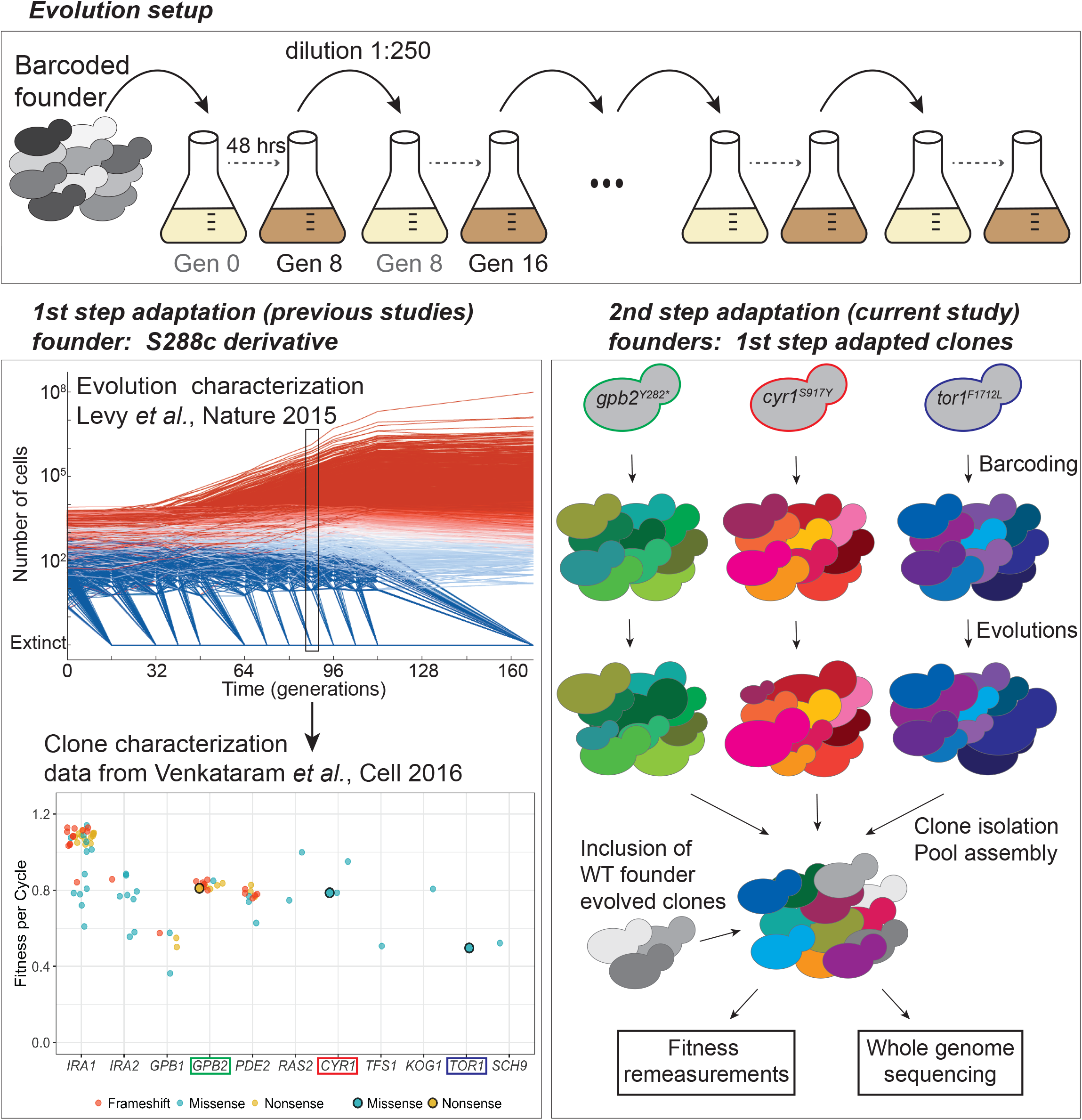
Experimental design. All evolutions were performed under identical conditions, including transfer and environmental conditions. The founders used in this study derived from adaptive clones isolated and characterized previously^41,4^. Each founder carries a single adaptive mutation (*gpb2^Y282*^, cyr1^S917Y^* and *tor1^F1712C^*), and was barcoded and propagated for 20 transfers. Adaptive clones were isolated and characterized via fitness re-measurements and whole genome sequencing.

### The Distribution of Fitness Effects (DFE) is compressed for the second adaptive step

We used lineage tracking data to estimate the distribution of beneficial fitness effects for each adapted ancestor from the 2^nd^-step evolutions and compare them to that of the wild-type ancestor from the 1^st^-step evolutions^41^ (datasets 1 and 2 in ^39^) (Fig. S2). Since the barcode sequencing depth was higher for the 1^st^-step evolutions (Table S3), we downsampled the 1^st^-step evolutions’ data to a depth comparable to that of the 2^nd^-step evolutions and calculated fitness and mutation rates (Fig. S5). Fitness inference remained similar upon downsampling (Fig. S5 A) and so did the mutation rate spectra for fitness above 4% (Fig. S5 B-C). We suspect that this discrepancy comes as a consequence of the faster adaptation of the wild type ancestor, resulting in very fit lineages dominating the population and neutral and lower fitness lineages thus being present at a low frequency. By down-sampling, we essentially limited our ability to detect lineages below ~4% fitness per generation, largely represented by diploids. Despite the lower barcode sequencing coverage, lineages with fitness <0.04 in the 2^nd^-step evolutions were readily detectable, in contrast to the 1^st^-step evolutions at comparable coverage (Fig. S5 C). Thus, high coverage lineage tracking data from ^39^ (datasets 1 and 2 without downsampling) are used for the 1^st^ step evolution, while low coverage lineage tracking data are used for the 2^nd^ step evolution.

Diminishing returns models of epistasis predict that the magnitude of fitness gains decreases as lineages approach a fitness optimum. Prior work has suggested that diminishing returns is at least partially due to decreased fitness gains as adaptive mutations occur along the same line^24,25,27,28,30^, however, the DFE beyond the first step has not been characterized. Our data allow us to directly compare the DFEs of two consecutive adaptation steps. We used the wild-type DFE for the first step and overlaid to the DFE of each of the adapted ancestors (Fig. 2). The density data were used to estimate mutation rates for diploidization and higher fitness mutations (Table 2). First, we observed a peak around fitness ~0.03-0.04 per generation across all genotypes. Our previous studies suggest that these peaks likely correspond to diploidization, shown to be adaptive in the wild-type background^4^. Follow-up ploidy assays and fitness re-measurements of individual clones confirmed the prevalence and fitness advantage of diploidization during the 2^nd^-step evolutions (see below for details). Additionally, 2^nd^-step evolutions manifest lower mutation rates over a wide fitness range beyond 0.04 per generation, and as the magnitude of fitness increases, the mutational fitness spectra decline at a faster rate compared to the 1^st^-step. Finally, adapted ancestors are devoid of very high fitness mutations compared to the wild-type ancestor, as demonstrated by the large difference in the mutation rates at fitness interval 0.07-0.12 and by the scarcity of lineages with fitness >0.12 for the adapted ancestors (Table 2). Overall, compared to the wild-type adaptation, the 2^nd^-step adaptation with *cyr1*, *gpb2* and *tor1* mutants as immediate ancestors not only have smaller magnitudes of fitness gains as expected based on previous studies^25^, but also consistently have lower beneficial mutation rates, which has not been previously characterized. This suggests that diminishing returns in our system is driven by both declining fitness gains and decreased beneficial mutation rates. Based on this change in the DFE, we hypothesize that the adaptive genetic bases during the 2^nd^-step evolution will differ, opening up the possibility that they are contingent on the first adaptive step.

**Figure 2.**
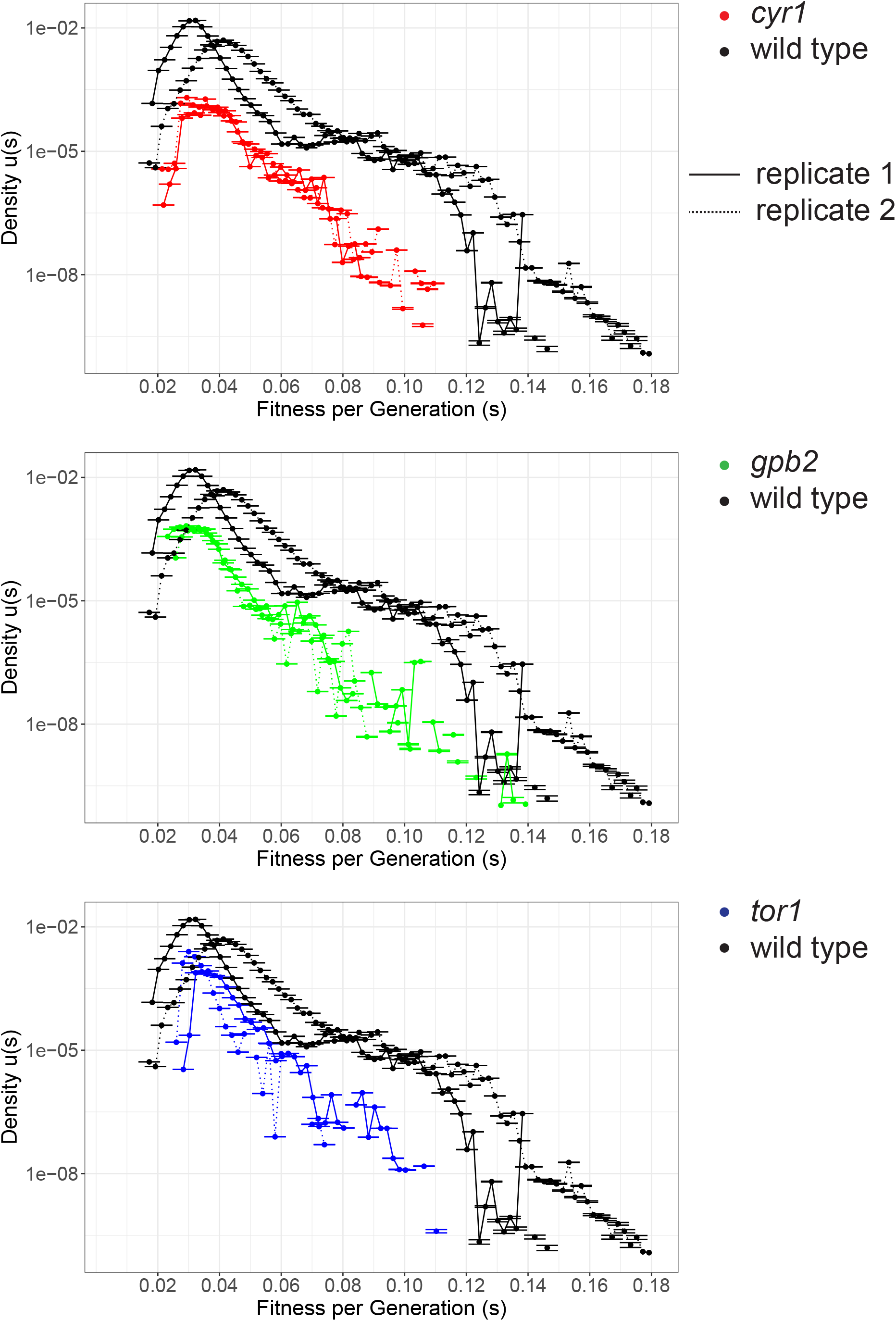
Mutation rates and fitness effects are smaller during the 2^nd^-step evolutions, as compared to the 1^st^-step. Mutation rates per fitness bin were calculated for all 2^nd^-step evolutions and compared to the 1^st^-step (wild type). The complete wild-type evolution datasets were taken into account for the calculation (see Fig. S5). Y-axis error bars are shown and min and max are annotated with a horizontal line (most errors are very small).

### Fitness increases of isolated adaptive clones from the 2^nd^-step evolutions tend to be smaller

Having analyzed the DFE from the lineage tracking data during the evolutions themselves, we followed up with a characterization of the distributions of fitness effects of individual clones isolated from each of the 2^nd^-step evolutions. Clones were isolated from a single timepoint from each evolution and their fitness effects were quantified against the wild-type ancestor under conditions identical to their evolutionary condition, by a bulk fitness assay. We directly compared the distribution of fitness effects between the 1^st^ and the 2^nd^-step evolutions, by including in our assays a set of isolated clones from the 1^st^-step evolutions^4^. Based on fitness and ploidy measurements, we classified isolated clones into four categories, consistent with the classification we used previously^4^. “Neutral haploids” are haploids with a similar fitness to their immediate ancestor. “Adaptive haploids” are haploids with a higher fitness compared to their immediate ancestor, presumably carrying adaptive mutation(s). “Pure diploids” are diploids without additional beneficial mutations. “High-fitness diploids” are diploids with a fitness significantly higher than the mean diploid fitness, and likely harbor beneficial mutation(s) besides diploidy.

Figure 3 shows the distributions of fitness effects per genotype, as calculated relative to the wild-type (panels to the left of the wild-type) and to their adapted ancestor (panels to the right of the wild-type). Isolated clones from the 1^st^-step evolution include the 2^nd^-step parental strains (corresponding points are annotated with larger dots in the wild-type panel, Fig. 3), whose fitness value is included in table S1 (under ‘Fitness evolved remeasurements’). Deviations from earlier estimates^4^ can be attributed to different population mean fitness resulting from inclusion of fitter strains in the pool. Neutral clones from the 2^nd^-step evolutions are expected to have fitness comparable to their respective parental strains from the 1^st^-step evolution, and that is the case for the *cyr1* and *gpb2* genotypes. However, neutral clones from the *tor1* genotype evolution have higher fitness than their unbarcoded ancestor, suggesting the possibility of the presence of mutation(s) that arose during the barcoding process. Overall, adapted clones from each of the three 2^nd^-step evolutions have further increased fitness compared to those from the 1^st^-step evolution (Fig. 3, fitness relative to the wild-type ancestor). However, the fitness increase of this 2^nd^ step is smaller than the fitness increase of the 1^st^ step (Fig. 3, fitness relative to the immediate ancestor), suggesting a slower adaptation rate, consistent with the data from the lineage tracking during the evolutions. In particular, during the 1^st^-step evolution, adaptive clones gain benefits up to ~0.18 per generation compared to their WT ancestor. During the 2^nd^ step evolutions, adaptive clones gain smaller fitness benefits compared to their immediate ancestors. The most fit clones gain benefits of ~0.09, ~0.10, and ~0.12 per generation compared to their *cyr1, gpb2*, and *tor1* ancestors, respectively. In agreement with diminishing returns, there is an anti-correlation between ancestor fitness and highest fitness evolved (Fig. S6, Pearson r = −0.95). We cross-validated our fitness estimates from the lineage tracking data from the evolutions and from the bulk competition assays, by plotting the estimates against each other (Fig. S7). Fitness values of lineages for which fitness was inferred from the evolution data approximately match the fitness values from the competition data. Discrepancies between the two datasets are expected to reflect cases where a single barcode represents more than a single genotype in the fitness inference from the lineage tracking data. The raw data for figures 3 and S7 are available in supplemental data, fitness dataset.

**Figure 3.**
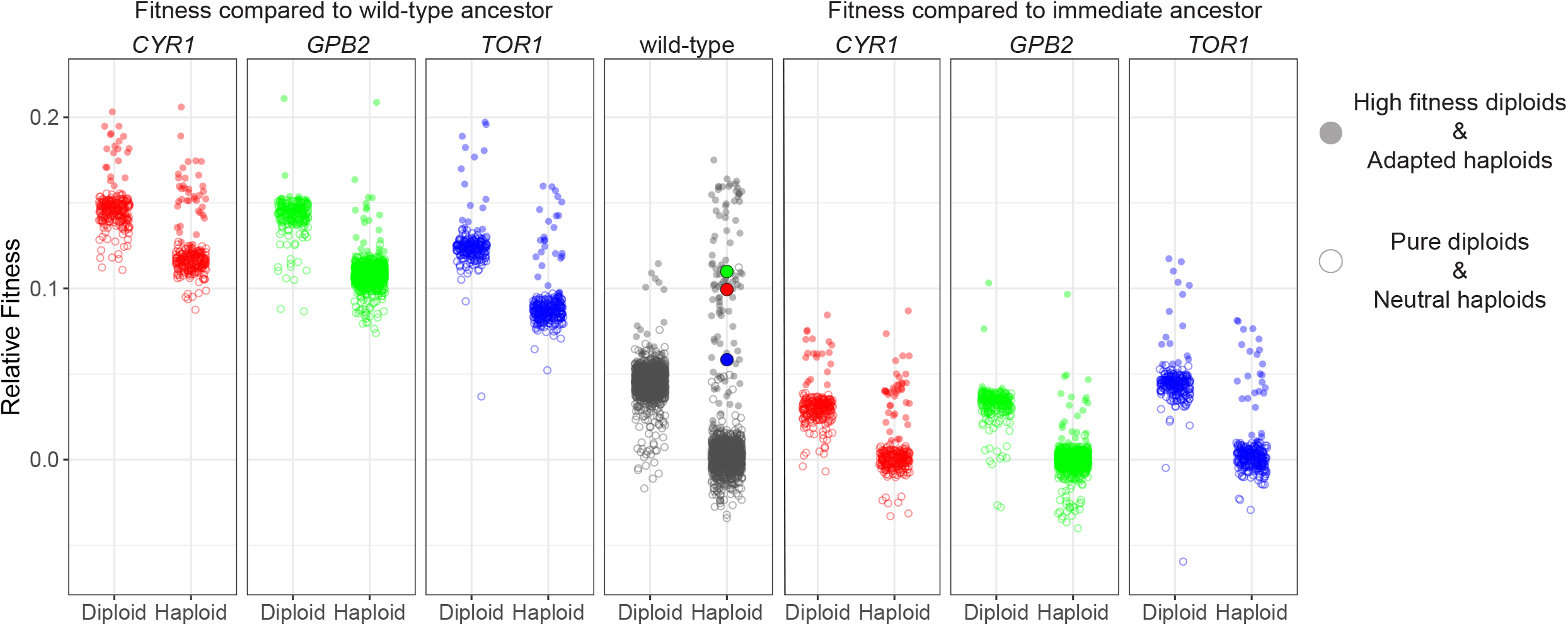
Distribution of fitness effects of 1^st^-step adapted clones following further adaptation. Fitness values of isolated clones are shown with respect to the wild-type ancestor (3 panels on the left) and with respect to their immediate ancestor (3 panels on the right). Fitness was measured in a pooled fashion from isolated clones of all immediate ancestor evolutions (including the wild-type), and arranged by ancestor. Haploids and diploids are shown in separate columns for each genotype. Clones with increased fitness within each group (high fitness diploids and adapted haploids) carry different annotation from pure diploids and neutral haploids. Larger dots in the ancestor cloud represent the adapted ancestors prior to barcoding and are color-coded to match the respective genotypes.

### Molecular targets of adaptation are contingent upon the founding genotype

To study the genetic basis of adaptation on the different genetic backgrounds, we performed whole genome sequencing on hundreds of adaptive clones isolated from the 2^nd^ step evolution. The genetic basis of the 1^st^ step evolution has been previously characterized^4^. Table 3 summarizes the molecular targets per founder and Fig. 4 shows their overlap in terms of genes and pathways. We observed similarities and differences in the mutational targets between the 1^st^- and 2^nd^-step evolutions and among the 2^nd^-step evolutions.

**Table 3:**
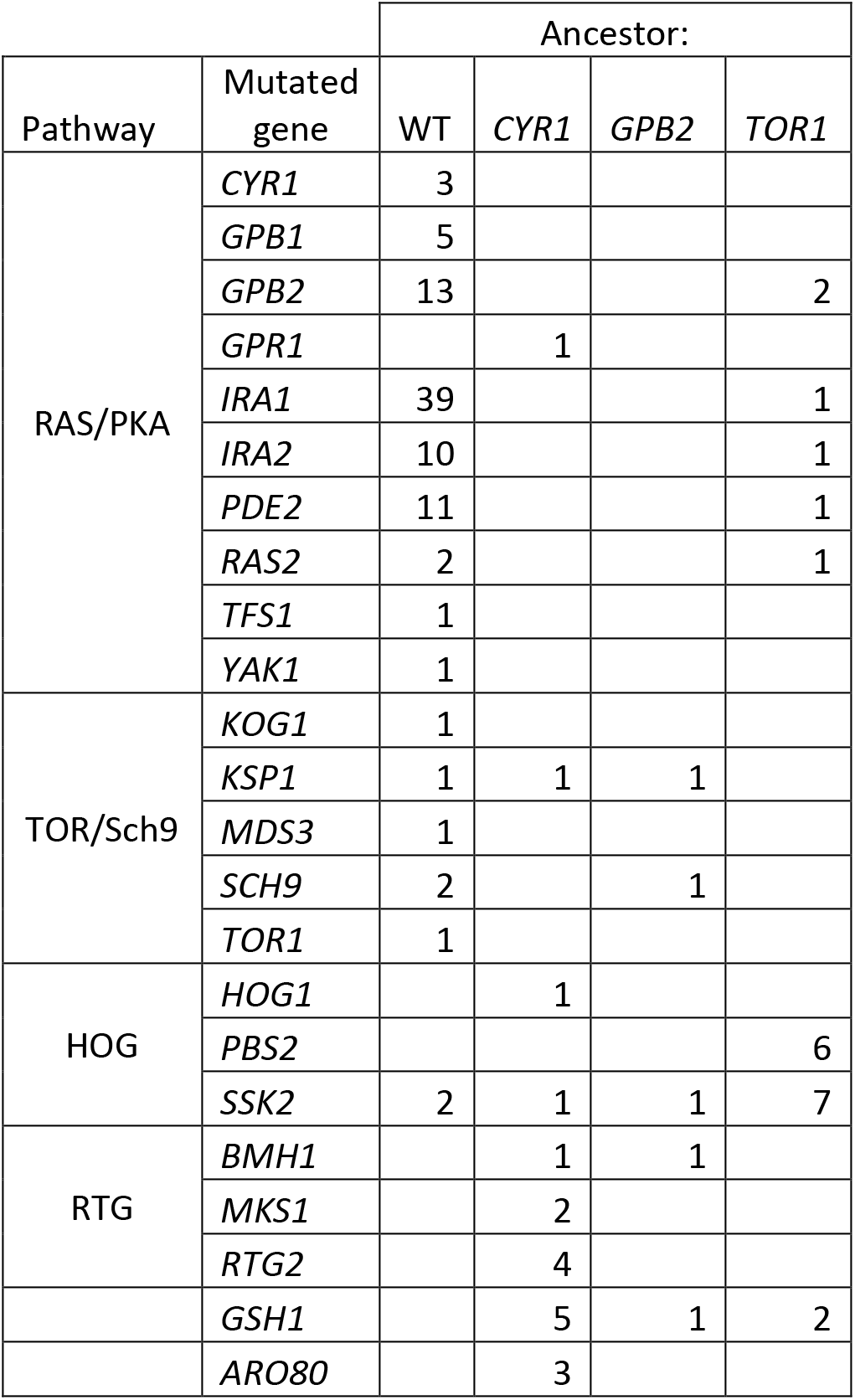
Genetic basis of adaptation per founder

**Figure 4.**
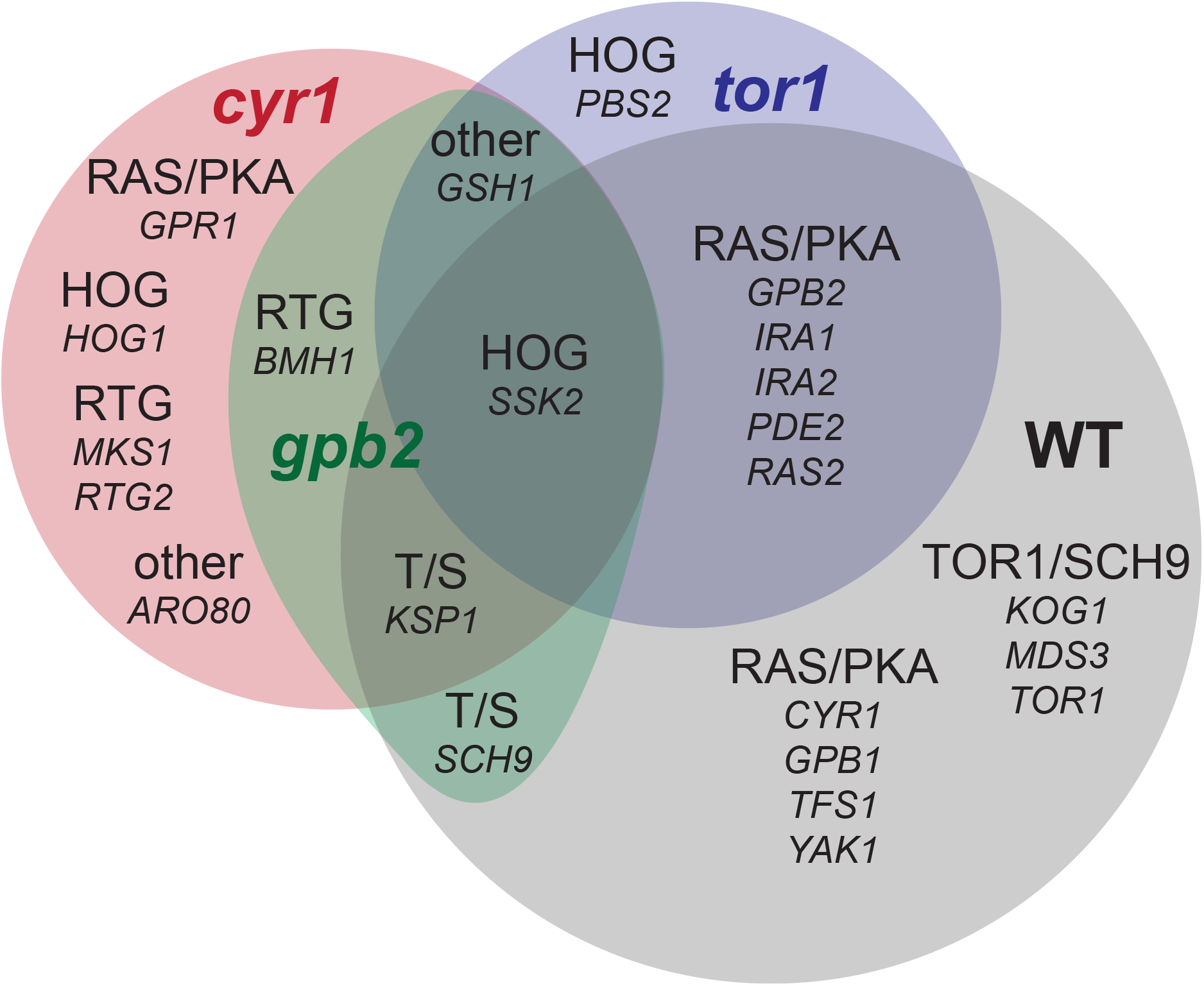
Overlap in the mutational recurrent targets among adaptive and wild-type ancestors. The pathway and the gene names are annotated in each area. For number of hits, see table 3. T/S stands for *TOR1*/*SCH9*.

Genes in the RAS/PKA and TOR/Sch9 pathways are the major adaptive targets during the 1^st^-step evolution and are also targets during the 2^nd^-step evolution. However, the *tor1* mutant is more likely to acquire adaptive mutations in the RAS/PKA pathway (6 out of 21 multi-hits in *tor1*, 1 out of 19 in *cyr1*, 0 out of 5 in *gpb2*), while *cyr1* and *gpb2* mutants are more likely to acquire mutations in the TOR/Sch9 pathway (1 out of 19 in *cyr1*, 2 out of 5 in *gpb2*, 0 out of 21 in *tor1*). The observation that double mutants on the RAS/PKA and TOR/Sch9 pathways are more fit than their corresponding single mutants and were selected for, whereas double mutants on the same pathway were not, suggests that the TOR and RAS/PKA pathways are not redundant in how they increase fitness, as has been previously shown^51^.

In contrast to the 1^st^-step targets of selection, stress response pathways were major targets of selection during the 2^nd^ evolutionary step. *GSH1* mutations were observed 8 times in total across all three genotypes of the 2^nd^-step evolution, yet no *GSH1* mutations were observed during the 1^st^-step evolution. Similarly, mutations affecting the retrograde (RTG) pathway were exclusively observed in the RAS/PKA mutant backgrounds (7 out of 19 in *cyr1*, 1 out of 5 in *gpb2*, 0 out of 21 in *tor1*), while HOG pathway mutants were observed predominantly in the *tor1* mutant background (13 out of 21 in *tor1* including pre-existing mutations, 2 out of 19 in *cyr1*, 1 out of 5 in *gpb2*) and *aro80* was also only observed in the *tor1* mutant background (Table 3, Fig. 4).

Finally, the predominant adaptive mutation type differs between 1^st^- and 2^nd^-step evolution, in terms of the consequence the mutations have on the encoded protein. In particular, adaptation via loss of function mutations is common during the 1^st^-step evolution, whereas the 2^nd^-step of adaptation often selects presumptive gain-of-function mutations. Adaptive changes that increase signaling in RAS/PKA and TOR/Sch9 pathways, can be achieved either by loss of function mutations in negative regulators, or, rarely, by presumptive gain of function in positive regulators. Specifically, 53 out of 95 causative mutations (56%) from the 1^st^ step evolution result in either a frameshift or stop-codon gained (nonsense), likely leading to the loss of function of the mutated gene. By contrast, only 14 out of 55 causative mutations (25%) from the 2^nd^ step evolution are frameshift or stop-codon gain mutations. In our calculations we included mutations that are presented in table 3 as common targets of selection, as well as mutations that occurred in the background of a stronger causal mutation candidate (not included in table 3, see Supplemental Data, adaptive targets and WGS datasets for detailed lists). The beneficial mutation types between the 1^st^- and 2^nd^-step evolutions are significantly different (chi-square p-value = 6E-4), which also indicates shifting beneficial mutation spectra during adaptation. It is also likely that if 2^nd^-step adaptation is more likely to result from gain-of-function mutations, then this may be partly responsible for decreased mutational target size, given that loss-of-function mutations occur more easily than gain-of-function mutations^52^.

### Negative epistasis may contribute to a narrower DFE

In theory, diminished fitness gains could result from allele pairs that interact negatively, Fitness(AB)<Fitness(A)+Fitness(B), and/or could be a reflection of the order in which adaptive mutations are selected (higher fitness mutations should be favored first). We estimated the extent of negative epistasis at the gene level. Availability of mutations affecting the same genes in both 1^st^- and 2^nd^-step evolutions, and of fitness effects of the implicated genotypes (single or double mutants), allows for a crude estimation of epistatic interactions among targets of selection. We considered the average fitness effects of alleles in two genes when they occur either singly (fitness averaged from all alleles of a gene, data from 1^st^-step evolutions) or together (fitness averaged from all genotypes that had the 2 genes mutated, data from 2^nd^-step evolutions) in the wild-type background, for genes where such data were available (Fig. 5). In all cases, 2^nd^-step adapted mutants are more fit than either of the 1^st^-step adapted mutants. The range of expected fitness for a genotype with both genes mutated, without epistasis between them (grey bar in Fig. 5), is represented by the 95% confidence interval of the sum of the mean fitness of mutants in each gene. Only *ksp1* in combination with either *cyr1* or *gpb2* was consistent with negative epistasis. The remaining combinations have fitness effects that do not deviate from the expectation of an additive model. Thus, these data only provide weak evidence for the hypothesis that negatively interacting alleles contribute to diminishing returns in our experiments.

**Figure 5.**
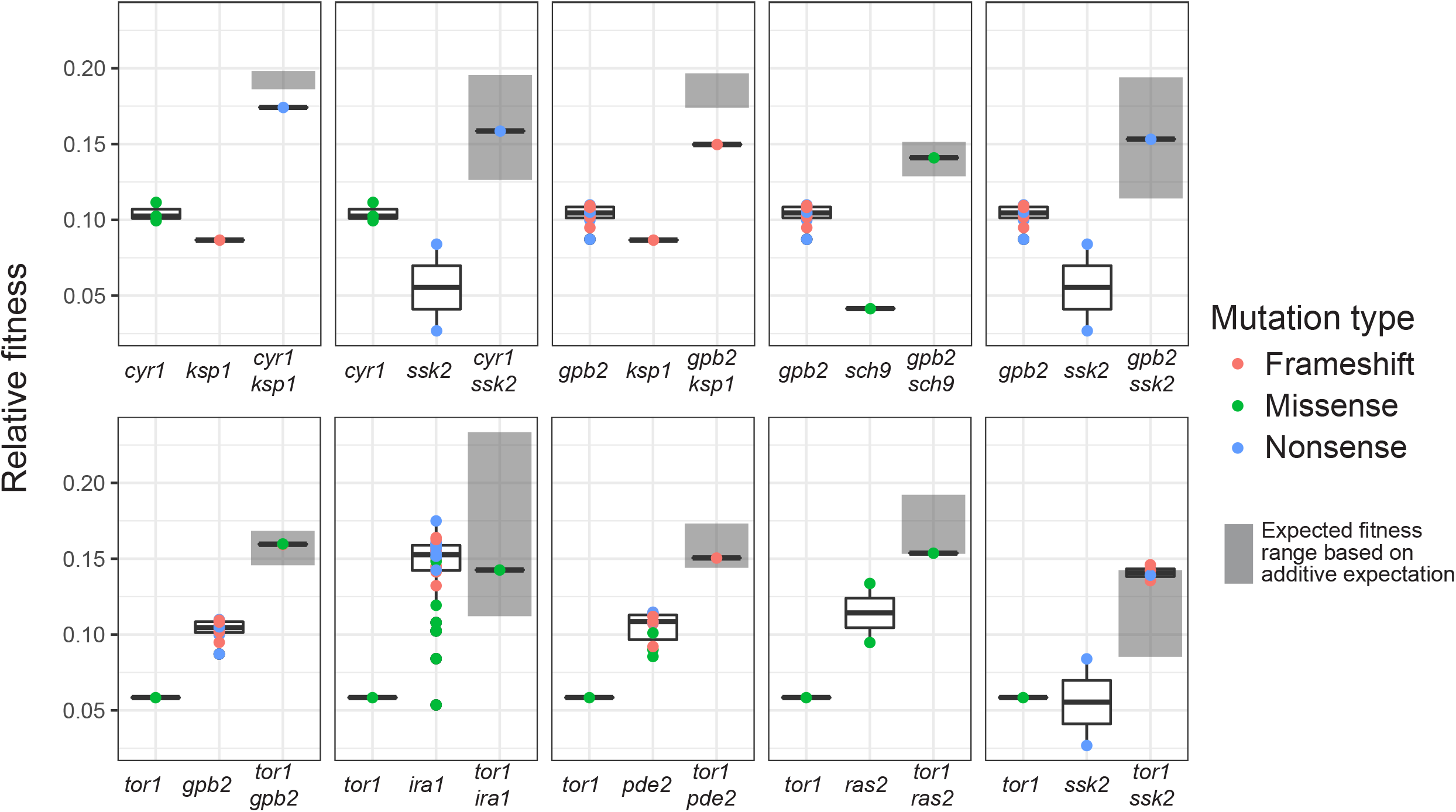
Pairwise interactions among targets of selection have additive or negative effects when combined. The additive fitness effects were estimated at gene-level for pairs of genes with adaptive alleles on the wild-type background (data from 1^st^-step evolution), and are annotated with grey bars, representing the 95% confidence interval of the sum of the mean fitness of mutants in each gene. Additive fitness effects are compared to the fitness effects of genotypes with adaptive alleles at both genes (data from 2^nd^-step evolutions), as a proxy for epistasis. The dot color of the double mutants refers to the second mutation shown in each graph. The first mutation matches the mutation type of the adapted ancestors (*cyr1* and *tor1* are missense and *gpb2* is a nonsense).

### Diploidization is adaptive across genotypes

Diploidization is a major adaptation strategy during evolution experiments founded with haploid yeast^4,53,54^. Previously, we demonstrated that during evolution of the wild-type ancestor, by generation 88, ~54% of the population was diploid. The majority of these diploids do not carry additional adaptive mutations and have a similar fitness advantage to one another over their wild-type ancestor (~0.045 per generation)^4^. Diploidization remained an adaptive strategy after acquisition of a first adaptive mutation across all genetic backgrounds. We directly estimated changes in the diploid fraction over evolutionary time based on benomyl sensitivity (Fig. S3). The assays suggested an increase in the fraction of diploid individuals over time (Fig. S3, Table S4). As a comparison, at generation 88 diploid individual fractions were on average 32%, 16%, 10% and 45% for the wild-type, *cyr1, gpb2* and *tor1* evolutions, based on assaying ~60-190 individuals per timepoint, for most timepoints. Interestingly, diploidization in the *tor1* background approached fixation in both replicates (96% and 88%) at generation 160, unlike the rest of the evolutions. That could be attributed to chance or may instead reflect either an otherwise comparatively weak adaptation potential for *tor1*, or an increased fitness for diploidy in a *tor1* background compared to the other backgrounds.

We used the fitness remeasurement data to calculate the fitness advantage of diploids in the context of different genetic backgrounds, including the wild-type ancestor. We observed two dominant groups of clones with distinct fitness (Fig. 3), which correspond to neutral haploids and pure diploids^4^. Pure diploids from wild-type, *cyr1, gpb2* and *tor1* genetic backgrounds have fitness advantages of ~0.045, 0.029, 0.031 and 0.043 per generation, respectively, relative to their immediate ancestors (Fig. 3). Although, these values are similar, there is an anti-correlation between ancestor fitness and diploidization fitness advantage (Fig. S6, Pearson r = −0.91), in concordance with the earlier observation that the highest fitness gains occur in the lowest fitness background. We also used the mutational fitness spectra to estimate diploidization mutation rates (Fig. 2). Diploidization mutation rates were assumed to correspond to the peaks of the mutation fitness spectra at the fitness interval ~3-4% per generation. The fitness values that correspond to these peaks along with their respective rates are shown in table 2. Average diploidization fitness across genotypes is comparable with the fitness estimates from the remeasurements data, albeit a little lower for the wild-type and *tor1* backgrounds. We consider the values from the fitness remeasurements more accurate and comparable to each other, since clones were competing in the same pool against a mostly wild-type population. The average across replicates mutation rates that correspond to these peaks are 2.03×10^−5^, 3.29×10^−7^, 1.19×10^−7^ and 3.35×10^−6^ for the wild-type, *cyr1*, *gpb2* and *tor1* ancestors, respectively. The value for the wild type is comparable to what has been previously reported on the diploidization rates in haploids during propagation^53^. The adapted genotypes display lower rates, matching the overall trend of lower mutation rates across all fitness intervals. Still, these data collectively suggest that diploidization remains a prevalent adaptation strategy after acquisition of a first adaptive mutation.

## Discussion

We characterized 2^nd^-step evolutions in terms of the DFE and adaptive mutational spectra and compared them to previously described 1^st^-step evolutions^4,41^. We used adapted clones, each carrying a single beneficial mutation compared to a common ancestor, as founders for the 2^nd^-step evolutions. We found that even a single step towards adaptation suffices to alter subsequent adaptation rates and adaptive mutational spectra.

Diminishing returns epistasis is apparent at multiple levels in our study, including the magnitude of fitness gains and the rate of adaptation. Both maximum and diploidization fitness gains anticorrelate with founder fitness (Fig. S6), in agreement with diminishing returns acting globally^25,30^. We also observed a lower mutation rate to modestly adaptive genotypes, as well as a depletion of high fitness events compared to the wild-type ancestor. These diminished adaptation rates are also supported by fitness remeasurements of individual clones from the 2^nd^-step evolution. Overall, these observations imply that both a smaller number of adaptive mutations *and* smaller fitness effects cause diminishing returns. Whole genome sequencing of adapted clones suggested that compared to the 1^st^ step of adaptation, the 2^nd^-step was more often the result of presumptive gain-of-function mutations. Such mutations are rarer^52^, and that may provide a potential explanation for why we observe a smaller adaptive target (equivalent to a lower adaptation rate) in our experiments.

In terms of their adaptive responses, wild-type and adaptive ancestors share common, converging, and diverging strategies. First, diploidization was a common and prevalent strategy among all genotypes. Second, adaptation through modifications of the RAS/PKA pathway was common between the *tor1* mutant and the wild-type ancestor, whereas adaptation through modifications of the TOR pathway was common among the RAS/PKA mutants (*cyr1* and *gpb2*) and the wild-type ancestor. This provides an example of convergent evolution, where genotypes with adaptive modifications in both pathways emerged from all three of the adapted ancestors. Combined with the fact that the same pathway was typically not mutated a second time (with one exception), this result is consistent with the idea that the Tor and Ras pathways are not redundant^51^. Sign epistasis between adaptive variants affecting the same pathway has been previously observed^22^, making it less likely for such double mutants to be selected. Finally, common adaptive responses during the 2^nd^-step evolution included mutations that potentially result in the upregulation of stress response pathways, which we did not observe in the 1^st^-step. Both TOR/Sch9 and RAS/PKA pathways, which were major targets during the 1^st^ step adaptation, regulate growth and stress responses responding to extracellular stimuli^51,54–57^. 1^st^-step adapted mutants had either RAS/PKA or TOR/Sch9 pathway upregulated^4,20^, which may decrease stress responses compared to their ancestor^56–66^. The adaptive basis of 2^nd^-step mutations may lie in the restoration of stress responses attenuated by overactive RAS/PKA or TOR/Sch9 pathways, as suggested by prior work^51,57–66^. In particular, all adapted ancestors evolved in this study acquired mutations in the *GSH1* gene, and evidence suggests that these mutations are gain-of-function, possibly resulting in an enhanced stress response. First, Gsh1p catalyzes the first and rate-limiting step of glutathione biosynthesis^67^, while no other mutations in the pathway were detected. Second, all *GSH1* mutations recovered were missense.

We also observed specific adaptation routes available to either the *tor1* mutant ancestor or the RAS/PKA mutant ancestors. Modifications of the retrograde pathway, including mutations in one positive (*RTG2*) and two negative (*BMH1* and *MKS1*) regulators^68^, were specific to the RAS/PKA mutants. In line with our observations on the glutathione biosynthesis pathway, all 4 mutations on *RTG2* were missense, while the 2 mutations on *BMH1* included a missense and a nonsense and the 2 mutations on *MKS1* included a missense and an upstream modification, suggesting that selection favors an enhanced retrograde flow pathway. Retrograde flow is negatively regulated by the TOR pathway^58,60,66^, providing a potential explanation as to why modifications of this particular pathway were not observed in *tor1* lineages: Modifications on retrograde flow, given an overactive TOR pathway, should be of larger effect, to overcome the additional load of the TOR-induced inhibition. Such large effect modifications may be rare, while *tor1* lineages are able to improve via different, more easily accessible routes, such as via modifications of the HOG pathway. Despite the common and converging adaptive responses, further adaptation also bears strong signatures of historical contingency, with mutational targets differing between wild type and adapted ancestors and between *tor1* and to RAS/PKA adapted ancestors. In particular, phenotypic relatedness dictates the degree of divergence, as shown by earlier studies^9^. We did not observe adaptive events contingent specifically on either the *cyr1* or the *gpb2* mutations, reflecting the phenotypic similarity of the *cyr1* and *gpb2* genotypes^4,9,20^. Nevertheless, that does not preclude that with a larger sample size we may have observed genotype-specific adaptive responses for *cyr1* and *gpb2*.

Our results provide clear evidence for the role of evolutionary history in shaping selection. This suggests that after acquisition of a single adaptive mutation the selective pressure a population experiences can change, even in the absence of environmental perturbation. The first adaptive change might be considered to be in direct response to the environmental condition, where adapted lineages modified their nutrient signaling pathways to respond to an environment that predictably undergoes glucose feast and famine^4,20^. Adaptive changes in pleiotropic genes (such as those that regulate nutrient signaling) may include non-adaptive or even maladaptive side effects. Thus, the set of second adaptive mutations may be constrained to adjust for pleiotropic consequences of the first, compensating for sub-optimal changes to the cellular network. This suggests that fine-tuning of the same pathway may be minimally beneficial in a majority of cases, compared to responses that adjust different pathways. Specific to our experiments, extensive literature (cited above) suggests that growth optimization comes with a cost in stress responses, and as a result, 2^nd^-step adaptation strategies targeting modification of stress responses may be contingent on the nature of the adaptive strategy caused by the 1^st^ step. This shift in adaptive strategy may underlie the observation that the 2^nd^ adaptive step was more often due to presumptive gain-of-function mutations.

## Supporting information

Adaptive Targets Data

Fitness Data

WGS data

Cyr1Evo1_CountsTimecourse

Cyr1Evo2_CountsTimecourse

Gpb2Evo1_CountsTimecourse

Gpb2Evo2_CountsTimecourse

Tor1Evo1_CountsTimecourse

Tor1Evo2_CountsTimecourse

## Figure legends

**Figure S1. The fitness advantage is a monogenic trait and co-segregates with the evolved variants.** Fitness advantage was measured against a fluorescent version of the initial ancestor, for each parental evolved clone and for 4 segregants derived from 2 consecutive back-crosses of the evolved clone with their ancestor. From each cross, one segregant with the evolved variant and one segregant with the ancestral variant were assayed. Fitness was measured in triplicates. Generations 8, 24 and 40 were used for fitness estimation of the cyr1 mutant and derivatives. Generations 0, 8, 24 and 40 were used for fitness estimations of the gpb2 and tor1 mutants and derivatives.

**Figure S2. Lineage trajectories over the course of the evolutions.** Lineage frequencies were estimated from time course barcode sequencing data. Data from the wild-type evolutions were subsampled from the counts in supplemental datasets 1 and 2 from^39^. **A.** Highlighted trajectories represent lineages with a fitness advantage and are colored by fitness. Grey trajectories represent neutral lineages, which are also shown in **B.** for clarity. In all cases 1000 neutral trajectories are shown.

**Figure S3. Frequency of adaptive individuals over the course of the evolutions.** The frequency was calculated using the barcode sequencing data.

**Figure S4. Diploid trajectories over the course of the evolutions.** Diploid ratio was estimated via a benomyl-based ploidy assay. Data of the wild-type evolution are the same as in ^4^, figure S7. Raw data are provided in table S4.

**Figure S5. Sequencing data downsampling does not affect the fitness calculation but affects the detection limit.** Barcode sequencing data from the 1^st^-step evolutions were downsampled to depths comparable to the data from the 2^nd^-step evolutions. **A.** Fitness effects were estimated using the complete and the downsampled datasets for both replicates^39^. **B-C.** Mutation rates per fitness bin as calculated using the complete (B) and the downsampled (C) datasets.

**Figure S6. Highest and diploidization fitness gains anticorrelate with ancestor fitness.** Fitness effects are expressed per generation. Pearson correlations equal −0.95 and −0.91, for highest and diploidization fitness effects, respectively.

**Figure S7. Fitness estimations from remeasurements of individual clones match the fitness values inferred from lineage tracking data of the evolutions.** Fitness values were matched based on barcode identity and ploidy characterization was based on a benomyl assay for individual clones.

## Acknowledgements

The authors wish to thank Lucas Herissant for help with FACS sorting and the benomyl assay, Jamie Blundell for help with data analysis, and Frank Rosenzweig and Sasha Levy for comments on the manuscript. The study was funded by NASA grant NNA15BB04A and NIH grant R35 GM131824 to GS.

## Notes

### Competing Interest Statement

The authors have declared no competing interest.

## REFERENCES

1. Gould, S. J. Wonderful Life: The Burgess Shale and the Nature of History. (W. W. Norton & Company, 1990).

2. Xie, K. T. et al. DNA fragility in the parallel evolution of pelvic reduction in stickleback fish. Science 363, 81–84 (2019).

3. Fisher, K. J., Kryazhimskiy, S. & Lang, G. I. Detecting genetic interactions using parallel evolution in experimental populations. Philosophical Transactions of the Royal Society B: Biological Sciences 374, 20180237 (2019).

4. Venkataram, S. et al. Development of a Comprehensive Genotype-to-Fitness Map of Adaptation-Driving Mutations in Yeast. Cell 166, 1585–1596.e22 (2016).

5. Sun, S., Coelho, M. A., Heitman, J. & Nowrousian, M. Convergent evolution of linked mating-type loci in basidiomycete fungi. bioRxiv 626911 (2019) doi:10.1101/626911.

6. Marcet-Houben, M. & Gabaldón, T. Evolutionary and functional patterns of shared gene neighbourhood in fungi. Nat Microbiol 1–10 (2019) doi:10.1038/s41564-019-0552-0.

7. Sackton, T. B. & Clark, N. Convergent evolution in the genomics era: new insights and directions. Philosophical Transactions of the Royal Society B: Biological Sciences 374, 20190102 (2019).

8. Deatherage, D. E., Kepner, J. L., Bennett, A. F., Lenski, R. E. & Barrick, J. E. Specificity of genome evolution in experimental populations of Escherichia coli evolved at different temperatures. PNAS 114, E1904–E1912 (2017).

9. Gac, M. L., Cooper, T. F., Cruveiller, S., Médigue, C. & Schneider, D. Evolutionary history and genetic parallelism affect correlated responses to evolution. Molecular Ecology http://onlinelibrary.wiley.com/doi/abs/10.1111/mec.12312 (2013) doi:10.1111/mec.12312.

10. Bailey, S. F., Rodrigue, N. & Kassen, R. The Effect of Selection Environment on the Probability of Parallel Evolution. Molecular Biology and Evolution 32, 1436–1448 (2015).

11. Conte, G. L., Arnegard, M. E., Peichel, C. L. & Schluter, D. The probability of genetic parallelism and convergence in natural populations. Proceedings of the Royal Society B: Biological Sciences 279, 5039–5047 (2012).

12. Stern, D. L. The genetic causes of convergent evolution. Nature Reviews Genetics 14, 751–764 (2013).

13. Blount, Z. D., Lenski, R. E. & Losos, J. B. Contingency and determinism in evolution: Replaying life’s tape. Science 362, eaam5979 (2018).

14. Moore, F. B.-G. & Woods, R. Tempo and constraint of adaptive evolution in Escherichia coli (Enterobacteriaceae, Enterobacteriales): TEMPO AND CONSTRAINT. Biological Journal of the Linnean Society 88, 403–411 (2006).

15. Ord, T. J. & Summers, T. C. Repeated evolution and the impact of evolutionary history on adaptation. BMC Evolutionary Biology 15, 137 (2015).

16. Echenique, J. I. R., Kryazhimskiy, S., Ba, A. N. N. & Desai, M. M. Modular epistasis and the compensatory evolution of gene deletion mutants. PLOS Genetics 15, e1007958 (2019).

17. Wytock, T. P. et al. Experimental evolution of diverse Escherichia coli metabolic mutants identifies genetic loci for convergent adaptation of growth rate. PLOS Genetics 14, e1007284 (2018).

18. Tenaillon, O. et al. The Molecular Diversity of Adaptive Convergence. Science 335, 457–461 (2012).

19. Wielgoss, S., Wolfensberger, R., Sun, L., Fiegna, F. & Velicer, G. J. Social genes are selection hotspots in kin groups of a soil microbe. Science 363, 1342–1345 (2019).

20. Li, Y. et al. Hidden Complexity of Yeast Adaptation under Simple Evolutionary Conditions. Current Biology 28, 515–525.e6 (2018).

21. Li, Y., Petrov, D. A. & Sherlock, G. Single nucleotide mapping of trait space reveals Pareto fronts that constrain adaptation. Nat Ecol Evol 3, 1539–1551 (2019).

22. Ono, J., Gerstein, A. C. & Otto, S. P. Widespread Genetic Incompatibilities between First-Step Mutations during Parallel Adaptation of Saccharomyces cerevisiae to a Common Environment. PLOS Biology 15, e1002591 (2017).

23. Hernando-Amado, S., Sanz-García, F. & Martínez, J. L. Antibiotic Resistance Evolution Is Contingent on the Quorum-Sensing Response in Pseudomonas aeruginosa. Molecular Biology and Evolution (2019) doi:10.1093/molbev/msz144.

24. Barrick, J. E., Kauth, M. R., Strelioff, C. C. & Lenski, R. E. Escherichia coli rpoB Mutants Have Increased Evolvability in Proportion to Their Fitness Defects. Mol Biol Evol 27, 1338–1347 (2010).

25. Wünsche, A. et al. Diminishing-returns epistasis decreases adaptability along an evolutionary trajectory. Nature Ecology & Evolution 1, 0061 (2017).

26. Gifford, D. R., Toll-Riera, M. & MacLean, R. C. Epistatic interactions between ancestral genotype and beneficial mutations shape evolvability in Pseudomonas aeruginosa. Evolution 70, 1659–1666 (2016).

27. MacLean, R. C., Perron, G. G. & Gardner, A. Diminishing Returns From Beneficial Mutations and Pervasive Epistasis Shape the Fitness Landscape for Rifampicin Resistance in Pseudomonas aeruginosa. Genetics 186, 1345–1354 (2010).

28. Moore, F. B.-G., Rozen, D. E. & Lenski, R. E. Pervasive compensatory adaptation in *Escherichia coli*. Proceedings of the Royal Society of London. Series B: Biological Sciences 267, 515–522 (2000).

29. Plucain, J. et al. Contrasting effects of historical contingency on phenotypic and genomic trajectories during a two-step evolution experiment with bacteria. BMC Evolutionary Biology 16, 86 (2016).

30. Kryazhimskiy, S., Rice, D. P., Jerison, E. R. & Desai, M. M. Global epistasis makes adaptation predictable despite sequence-level stochasticity. Science 344, 1519–1522 (2014).

31. Kvitek, D. J. & Sherlock, G. Reciprocal Sign Epistasis between Frequently Experimentally Evolved Adaptive Mutations Causes a Rugged Fitness Landscape. PLOS Genetics 7, e1002056 (2011).

32. Chiotti, K. E. et al. The Valley-of-Death: Reciprocal sign epistasis constrains adaptive trajectories in a constant, nutrient limiting environment. Genomics 104, 431–437 (2014).

33. Tenaillon, O. et al. Tempo and mode of genome evolution in a 50,000-generation experiment. Nature 536, 165–170 (2016).

34. Kao, K. C. & Sherlock, G. Molecular characterization of clonal interference during adaptive evolution in asexual populations of *Saccharomyces cerevisiae*. Nature Genetics 40, 1499–1504 (2008).

35. Good, B. H., McDonald, M. J., Barrick, J. E., Lenski, R. E. & Desai, M. M. The dynamics of molecular evolution over 60,000 generations. Nature 551, 45–50 (2017).

36. Long, A., Liti, G., Luptak, A. & Tenaillon, O. Elucidating the molecular architecture of adaptation via evolve and resequence experiments. Nature Reviews Genetics 16, 567–582 (2015).

37. Lang, G. I. et al. Pervasive genetic hitchhiking and clonal interference in forty evolving yeast populations. Nature 500, 571–574 (2013).

38. Gerrish, P. J. & Lenski, R. E. The fate of competing beneficial mutations in an asexual population. Genetica 102, 127 (1998).

39. Blundell, J. R. et al. The dynamics of adaptive genetic diversity during the early stages of clonal evolution. Nature Ecology & Evolution 3, 293 (2019).

40. Ba, A. N. N. et al. High-resolution lineage tracking reveals travelling wave of adaptation in laboratory yeast. Nature 575, 494–499 (2019).

41. Levy, S. F. et al. Quantitative evolutionary dynamics using high-resolution lineage tracking. Nature 519, 181–186 (2015).

42. Daniel Gietz, R. & Woods, R. A. Transformation of yeast by lithium acetate/single-stranded carrier DNA/polyethylene glycol method. in Methods in Enzymology (eds. Guthrie, C. & Fink, G. R.) vol. 350 87–96 (Academic Press 2002).

43. Verduyn, C., Postma, E., Scheffers, W. A. & Van Dijken, J. P. Effect of benzoic acid on metabolic fluxes in yeasts: A continuous-culture study on the regulation of respiration and alcoholic fermentation. Yeast 8, 501–517 (1992).

44. Jaffe, M., Sherlock, G. & Levy, S. F. iSeq: A New Double-Barcode Method for Detecting Dynamic Genetic Interactions in Yeast. G3: Genes, Genomes, Genetics 7, 143–153 (2017).

45. Zhao, L., Liu, Z., Levy, S. F. & Wu, S. Bartender: a fast and accurate clustering algorithm to count barcode reads. Bioinformatics 34, 739–747 (2018).

46. Baym, M., Shaket, L., Anzai, I. A., Adesina, O. & Barstow, B. Rapid construction of a whole-genome transposon insertion collection for *Shewanella oneidensis* by Knockout Sudoku. Nature Communications 7, 13270 (2016).

47. Baym, M. et al. Inexpensive Multiplexed Library Preparation for Megabase-Sized Genomes. PLOS ONE 10, e0128036 (2015).

48. Martin, M. Cutadapt removes adapter sequences from high-throughput sequencing reads. EMBnet.journal 17, 10–12 (2011).

49. Li, H. & Durbin, R. Fast and accurate short read alignment with Burrows-Wheeler transform. Bioinformatics 25, 1754–1760 (2009).

50. Cingolani, P. et al. A program for annotating and predicting the effects of single nucleotide polymorphisms, SnpEff: SNPs in the genome of Drosophila melanogaster strain w ^1118^; iso-2; iso-3. Fly 6, 80–92 (2012).

51. Kunkel, J., Luo, X. & Capaldi, A. P. Integrated TORC1 and PKA signaling control the temporal activation of glucose-induced gene expression in yeast. Nat Commun 10, 1–11 (2019).

52. Zörgö, E. et al. Life History Shapes Trait Heredity by Accumulation of Loss-of-Function Alleles in Yeast. Mol Biol Evol 29, 1781–1789 (2012).

53. Harari, Y., Ram, Y., Rappoport, N., Hadany, L. & Kupiec, M. Spontaneous Changes in Ploidy Are Common in Yeast. Current Biology 28, 825–835.e4 (2018).

54. Santangelo, G. M. Glucose Signaling in Saccharomyces cerevisiae. Microbiol. Mol. Biol. Rev. 70, 253–282 (2006).

55. Tamaki, H. Glucose-stimulated cAMP-protein kinase a pathway in yeast Saccharomyces cerevisiae. Journal of Bioscience and Bioengineering 104, 245–250 (2007).

56. Campos, S. E. & DeLuna, A. Functional genomics of dietary restriction and longevity in yeast. Mechanisms of Ageing and Development 179, 36–43 (2019).

57. Liu, Y., Yang, F., Li, S., Dai, J. & Deng, H. Glutaredoxin Deletion Shortens Chronological Life Span in Saccharomyces cerevisiae via ROS-Mediated Ras/PKA Activation. J. Proteome Res. 17, 2318–2327 (2018).

58. Conrad, M. et al. Nutrient sensing and signaling in the yeast Saccharomyces cerevisiae. FEMS Microbiol Rev 38, 254–299 (2014).

59. Ahmed, K., Carter, D. E. & Lajoie, P. Hyperactive TORC1 sensitizes yeast cells to endoplasmic reticulum stress by compromising cell wall integrity. FEBS Letters 593, 1957–1973 (2019).

60. Rutherford, J. C., Bahn, Y.-S., van den Berg, B., Heitman, J. & Xue, C. Nutrient and Stress Sensing in Pathogenic Yeasts. Front. Microbiol. 10, (2019).

61. Galdieri, L., Mehrotra, S., Yu, S. & Vancura, A. Transcriptional Regulation in Yeast during Diauxic Shift and Stationary Phase. OMICS: A Journal of Integrative Biology 14, 629–638 (2010).

62. Jamieson, D. J. Oxidative stress responses of the yeast Saccharomyces cerevisiae. Yeast 14, 1511–1527 (1998).

63. Charizanis, C., Juhnke, H., Krems, B. & Entian, K.-D. The oxidative stress response mediated via Pos9/Skn7 is negatively regulated by the Ras/PKA pathway in Saccharomyces cerevisiae. Mol Gen Genet 261, 740–752 (1999).

64. Iida, H. Multistress resistance of Saccharomyces cerevisiae is generated by insertion of retrotransposon Ty into the 5’ coding region of the adenylate cyclase gene. Molecular and Cellular Biology 8, 5555–5560 (1988).

65. Anand, A. et al. Adaptive evolution reveals a tradeoff between growth rate and oxidative stress during naphthoquinone-based aerobic respiration. PNAS 116, 25287–25292 (2019).

66. Rødkær, S. V. & Færgeman, N. J. Glucose- and nitrogen sensing and regulatory mechanisms in Saccharomyces cerevisiae. FEMS Yeast Res 14, 683–696 (2014).

67. Meister, A. Glutathione Metabolism and Its Selective Modification. The Journal of Biological Chemistry 263, 17205–17208 (1988).

