## Supplementary figures and images for "Changes in the distribution of fitness effects and adaptive mutational spectra following a single first step towards adaptation"

### Supplementary Figures

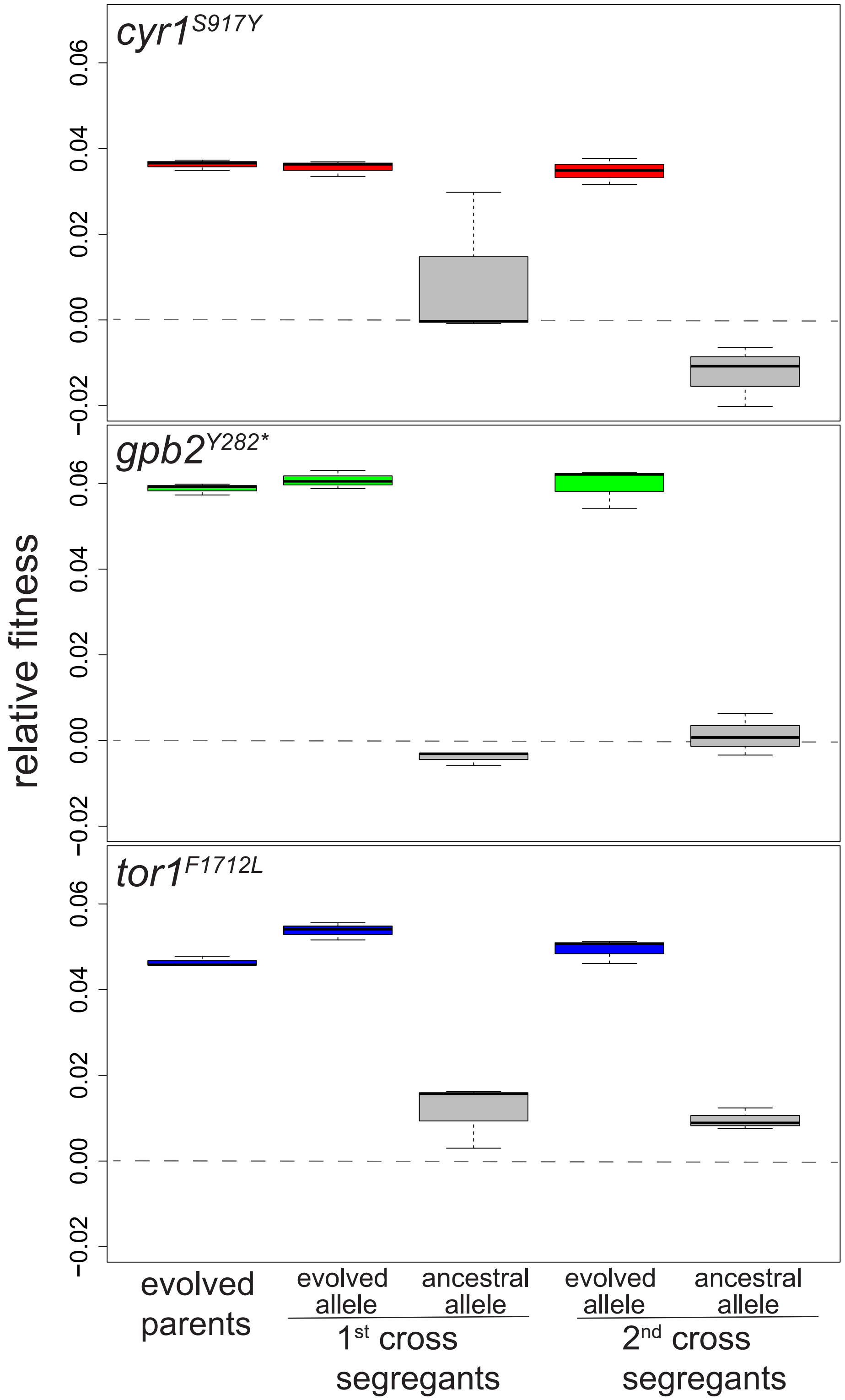

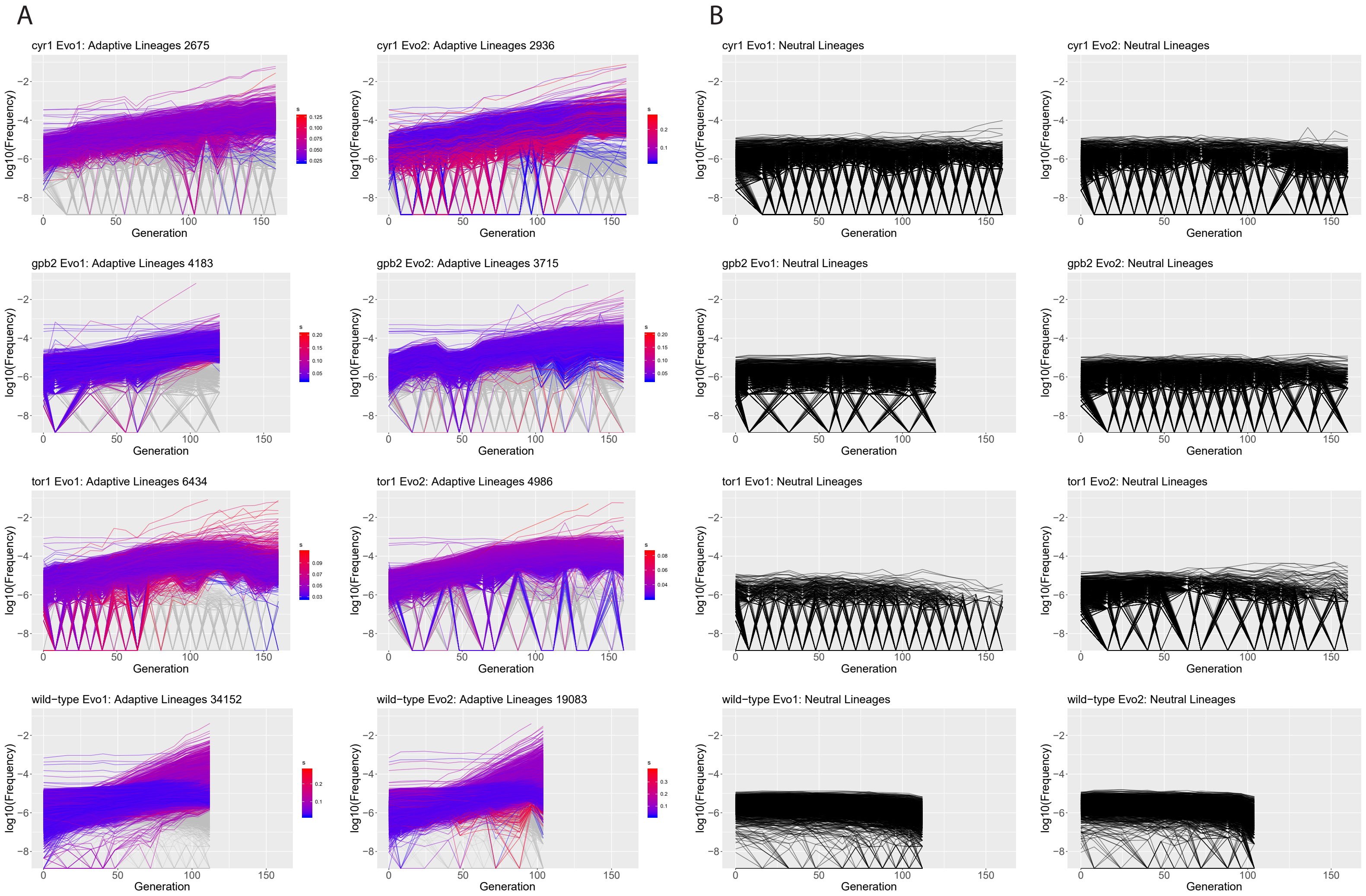

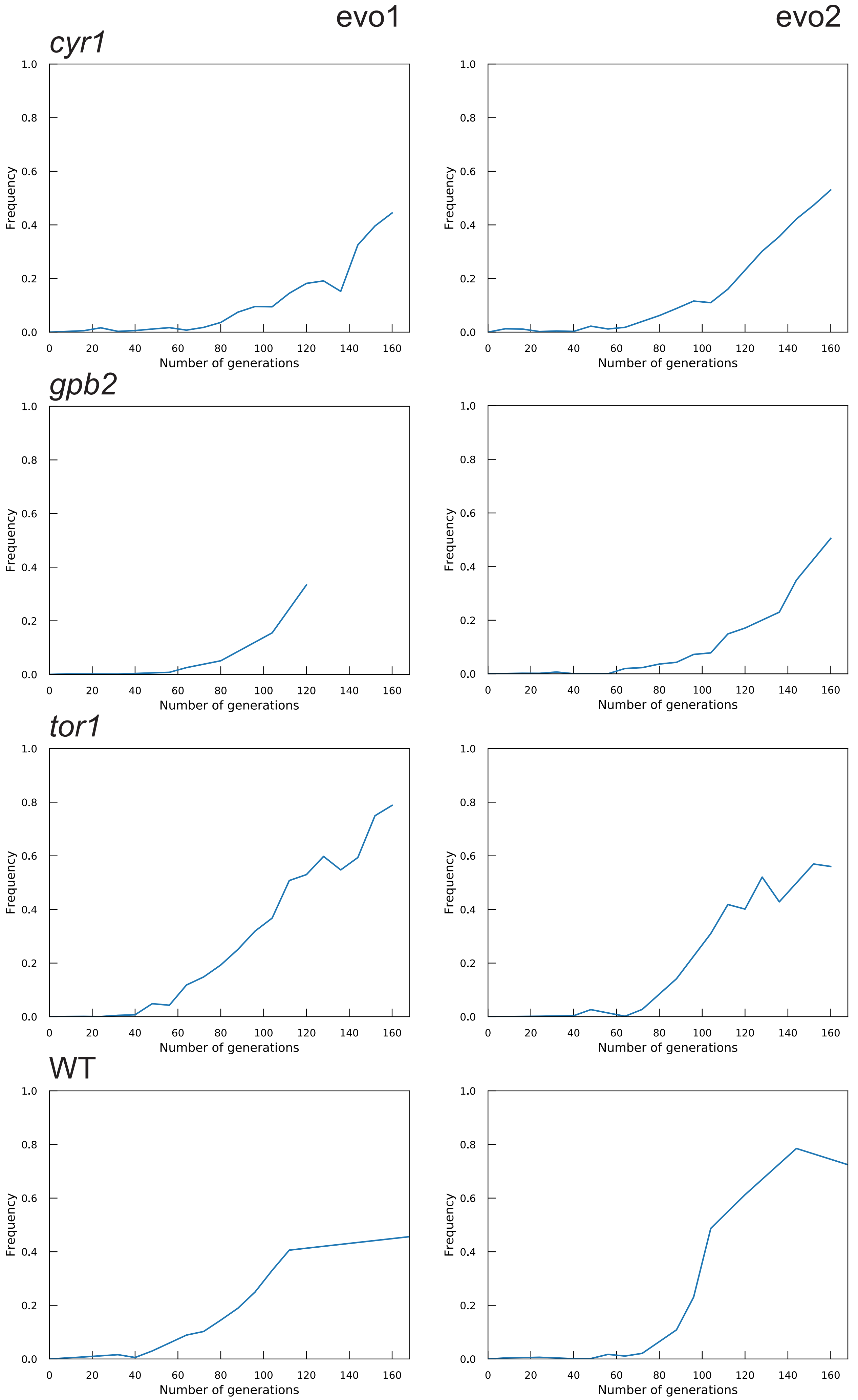

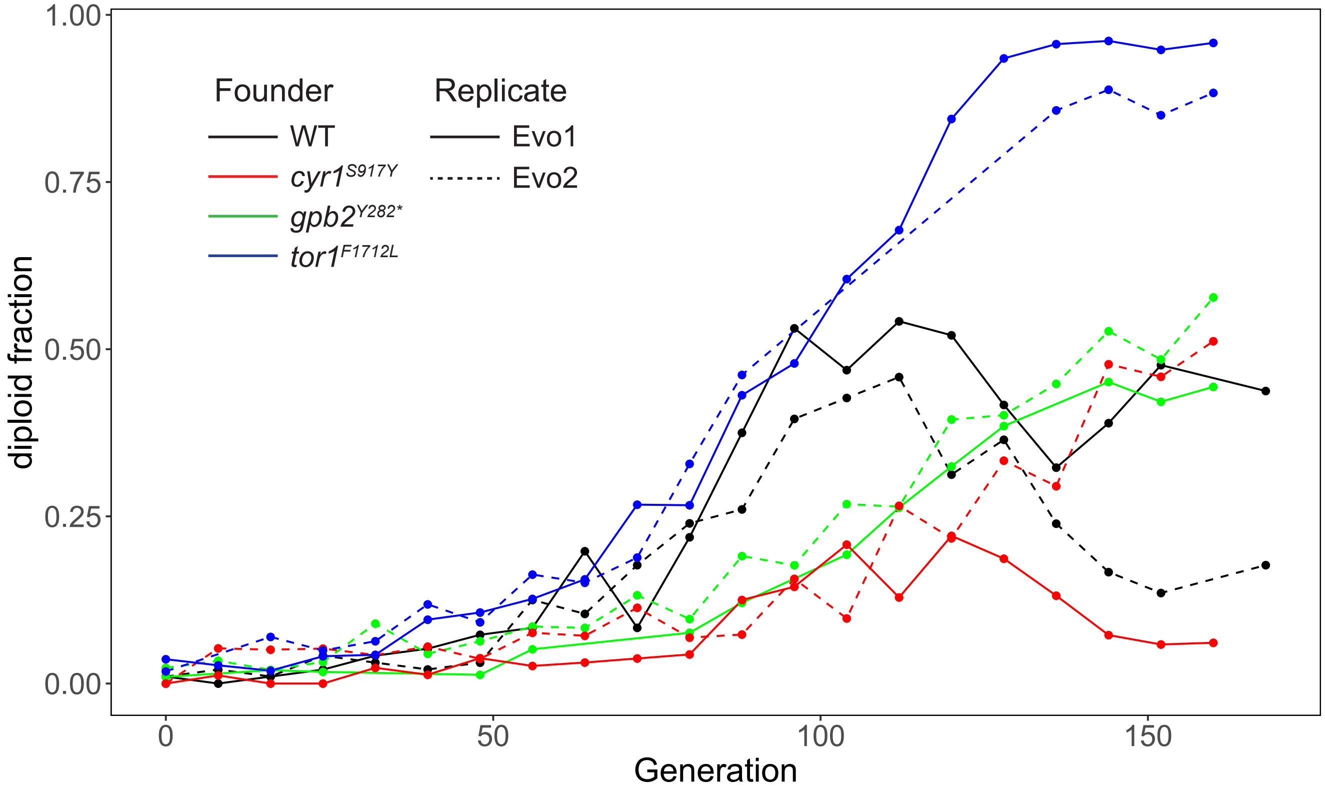

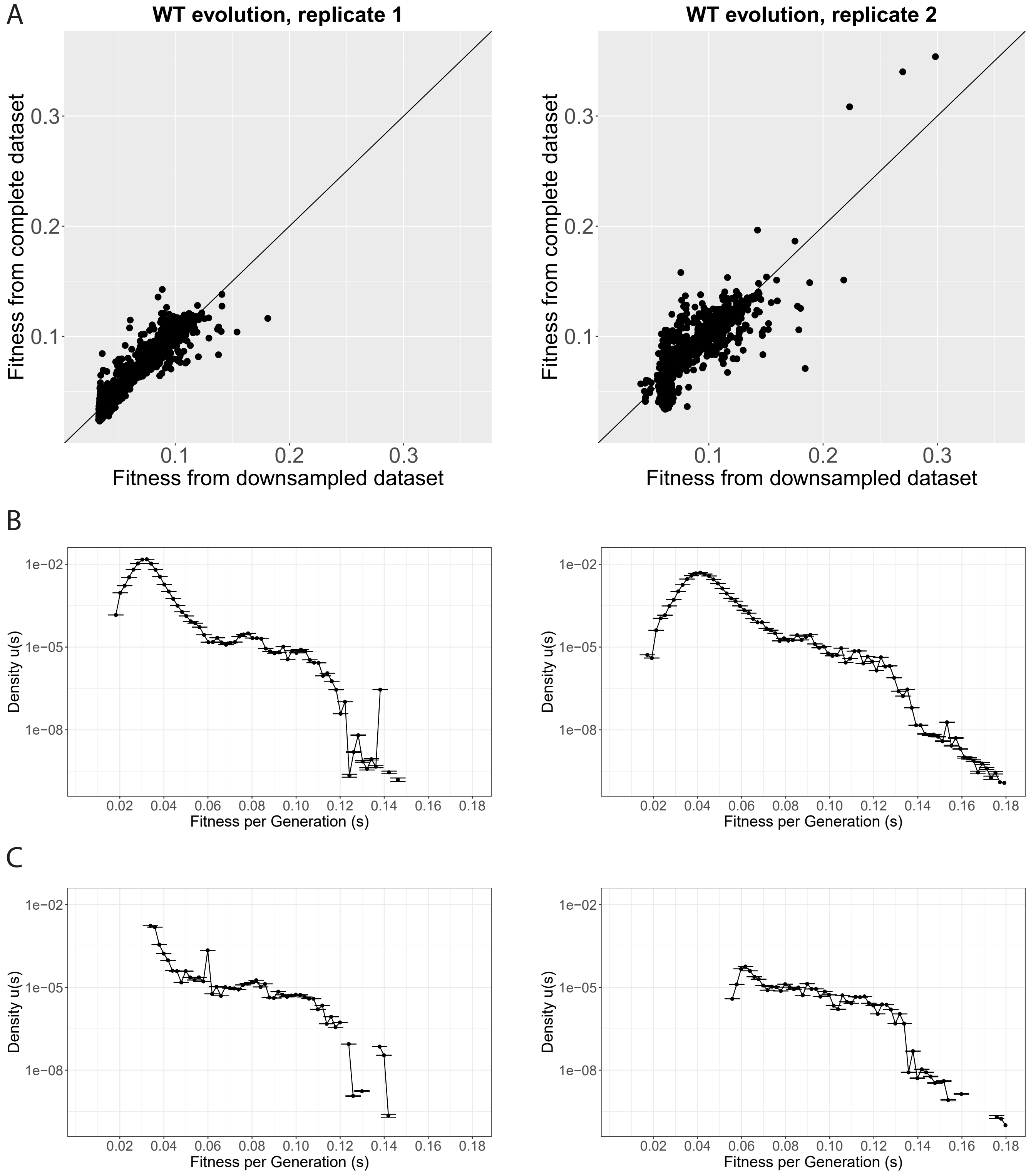

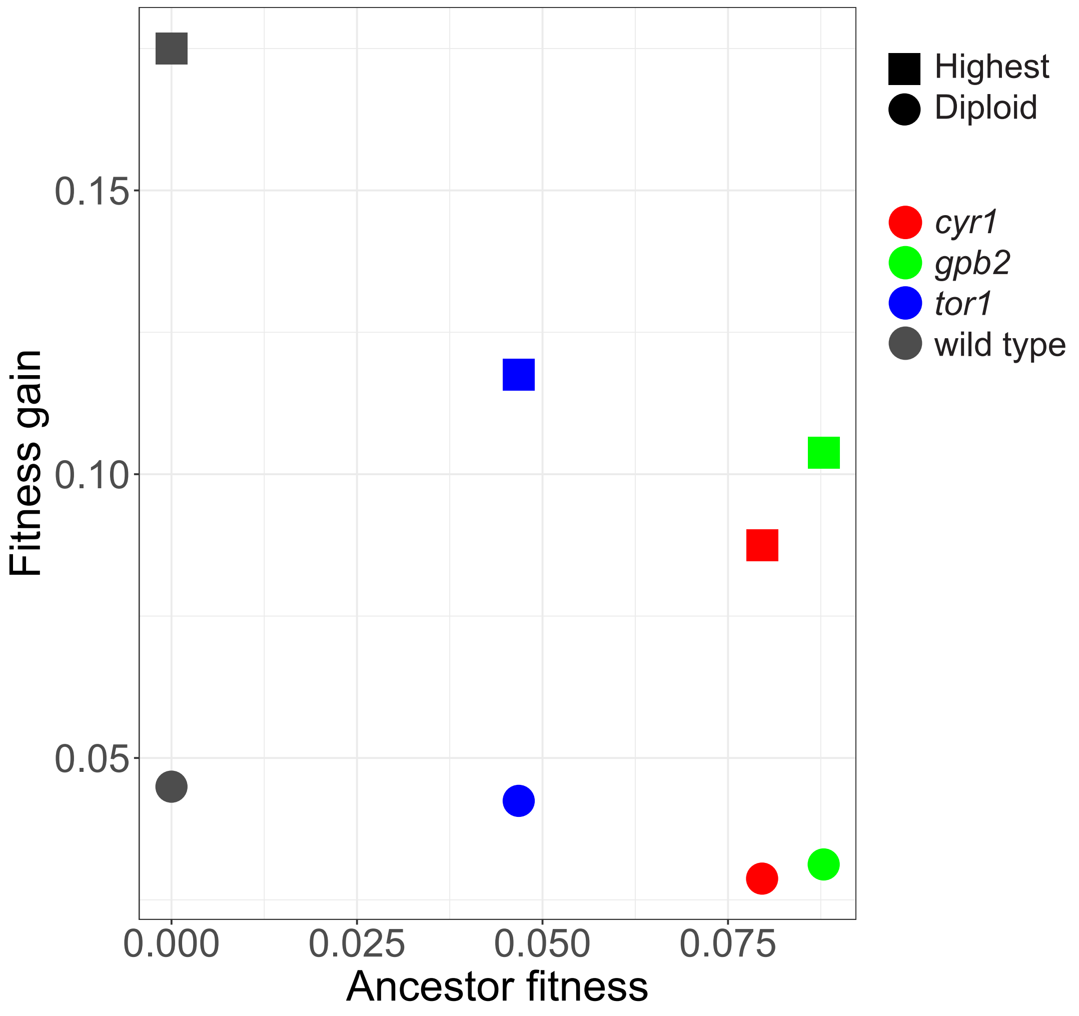

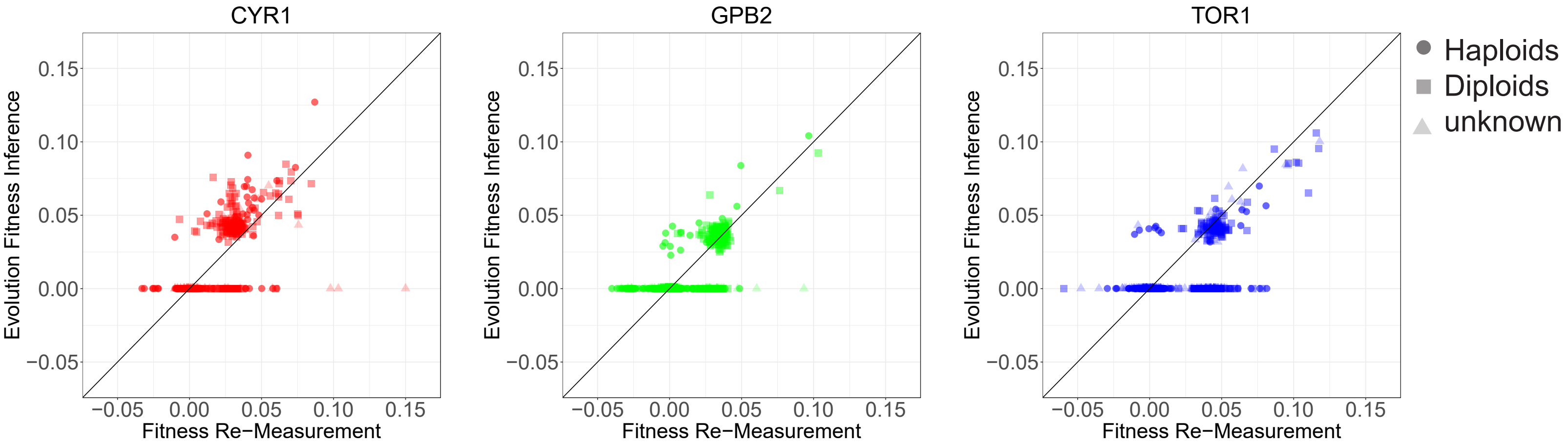
